# Endo-lysosomal assembly variations among Human Leukocyte Antigen class I (HLA-I) allotypes

**DOI:** 10.1101/2022.03.31.485520

**Authors:** Eli Olson, Theadora Ceccarelli, Malini Raghavan

## Abstract

The extreme polymorphisms of HLA-I proteins enable the presentation of diverse peptides to cytotoxic T lymphocytes (CTL). The canonical endoplasmic reticulum (ER) HLA-I assembly pathway enables presentation of cytosolic peptides, but effective intracellular surveillance requires multi-compartmental antigen sampling. Endo-lysosomes are generally sites of HLA class II assembly, but human monocytes and monocyte-derived dendritic cells (moDCs) also contain significant reserves of endo-lysosomal HLA-I molecules. We hypothesized variable influences of HLA-I polymorphisms upon outcomes of endo-lysosomal trafficking, as the stabilities and peptide occupancies of cell surface HLA-I are variable. For example, in moDCs, compared with HLA-B*08:01, HLA-B*35:01 displays reduced cell-surface stability and greater receptivity to exogenous peptide. Perturbations of endo-lysosomal pH negatively affect the surface expression of moDC HLA-B*35:01 but not HLA-B*08:01, causing HLA-B*35:01 accumulation in LAMP1^+^ compartments. These findings reveal the intersection of the vacuolar cross-presentation pathway with a constitutive assembly pathway for HLA-B*35:01. Notably, cross-presentation of epitopes derived from two soluble antigens was also more efficient for B*35:01 compared to B*08:01, even when matched for T cell response sensitivity, and more affected by cathepsin inhibition. Thus, HLA-I polymorphisms dictate the degree of endo-lysosomal assembly, which can supplement ER assembly for constitutive HLA-I expression and increase the efficiency of cross-presentation.

## INTRODUCTION

The major histocompatibility complex class I (MHC-I) is responsible for providing nearly all cells in the body with the capability for intracellular immune surveillance. This is accomplished by presenting intracellular peptides on the cell surface, where they are recognized by CD8^+^ T cells (Bjorkman and Parham, 1990). CD8^+^ T cells that have been properly selected in the thymus do not react to self-derived peptides but may recognize and activate in response to foreign antigen (Klein et al., 2014). Assembly of MHC-I complexes with peptides typically occurs in the endoplasmic reticulum (ER), where nascent MHC-I heavy chains are synthesized and associate with the invariant β _2_m light chain, followed by binding to a complex of chaperones known as the peptide loading complex (PLC) (Blum et al., 2013). MHC-I heavy chain/light chain dimers are stabilized by the PLC and brought into close association with the transporter associated with antigen processing (TAP), a dimeric peptide transporter that brings cytosolically processed peptides into the ER for peptide loading. Upon successful assembly, MHC-I complexes are sufficiently stabilized so that they can release from the PLC and traffic to the cell surface (Blum et al., 2013, Raghavan and Geng, 2015). Assembly is a highly coordinated and regulated process that is often targeted by viruses and cancers to escape immune surveillance (Hansen and Bouvier, 2009, Verweij et al., 2015, Leone et al., 2013), thus highlighting the importance of efficient assembly and surface MHC-I expression.

Human MHC-I (human leukocyte antigen class I (HLA-I)) heavy chains are encoded by three highly polymorphic genes: HLA-A, HLA-B, and HLA-C, with HLA-B being the most polymorphic of the three. The high polymorphisms enable the presentation of diverse antigens to CD8^+^ T cells (Falk et al., 1991, Sarkizova et al., 2020). The polymorphisms also result in divergent assembly and expression variations among allotypes, causing both stability (Rizvi et al., 2014, Geng et al., 2018a) and peptide repertoire differences (Falk et al., 1991, Sarkizova et al., 2020). HLA-B peptide preferences have a strong effect on the surface expression levels in primary lymphocytes; for example, allotypes that prefer peptides with a proline at position 2 (P2P peptides) are expressed at relatively low levels, display faster dissociation rates, and have more peptide receptive forms on the surface of T lymphocytes (Yarzabek et al., 2018). Most P_2_P-preferring allotypes fall into the B7 supertype and include related allotypes such as B*35:01 and B*07:02 (Sidney et al., 2008). Interestingly, the transporter associated with antigen processing (TAP) disfavors transport of P_2_P peptides (Uebel et al., 1997), creating a mismatch in peptide binding preferences and potentially a depleted supply of P_2_P peptides in the ER. Also of note, B7 members such as B*35:01 and B*07:02 assemble and express at the cell surface at relatively high levels in the absence of TAP (Geng et al., 2018b), indicating a predisposition for non-canonical assembly pathways. Additionally, several HLA-I allotypes, including some B7 supertype members, can assemble independently of tapasin (Peh et al., 1998, Williams et al., 2002, Rizvi et al., 2014), a key PLC component (Blum et al., 2013). Tapasin is known to edit the HLA-I peptide repertoire towards high affinity sequences (Rizvi and Raghavan, 2006, Wearsch and Cresswell, 2007, Chen and Bouvier, 2007) and tapasin-deficient cells have lower cell surface stability (Garbi et al., 2003) than their wild type counterparts. Allotypes that can assemble independently of tapasin may generally contain suboptimal peptide repertoires, resulting in complexes that are less stable (more rapidly endocytosed (Zagorac et al., 2012)) and more peptide receptive on the cell surface (a phenotype induced by suboptimal peptide loading (Ljunggren et al., 1990, Schumacher et al., 1990)). Indeed, these characteristics are measurable for B*35:01 and B*07:02 in primary lymphocytes (Yarzabek et al., 2018).

Following expression on the cell surface via the secretory pathway, MHC-I molecules are endocytosed, sorted, and recycled to the cell surface or trafficked to lysosomes for degradation; these are well-studied processes (Montealegre and van Endert, 2018). The relative activities of these pathways can be variable between cell types; for example, immature DCs are generally more endocytic compared to mature DCs (Trombetta and Mellman, 2005). In addition to ER assembly characteristics, the extent of endocytosis, recycling, lysosomal targeting, and lysosomal degradative capacity can each influence cell surface expression levels and half-lives of HLA-I molecules in a cell-type dependent manner. For example, immature Langerhans cells have substantial endosomal stores of MHC-I which re-distribute to the cell surface and become more stable upon maturation (MacAry et al., 2001), indicative of a shift in trafficking and MHC-I assembly. Conditions for peptide binding within endosomal and phagosomal compartments are quite different compared to canonical assembly in the ER due to lower compartmental pH within endo-lysosomes and the lack of ER chaperones and PLC components. Based on the marked variation in ER assembly characteristics of HLA-I allotypes (Rizvi et al., 2014, Geng et al., 2018b, Peh et al., 1998, Williams et al., 2002), we hypothesized that there would be differences in the outcomes of endocytic and lysosomal trafficking in both allotype and cell type dependent manners. HLA-I allotypes such as B*35:01 that undergo suboptimal ER assembly and can escape PLC-mediated quality control (Yarzabek et al., 2018, Rizvi et al., 2014, Geng et al., 2018b) may be particularly well suited for assembly or survival within endo-lysosomal compartments in cells that maintain significant pools of endo-lysosomal HLA-I.

Dendritic cells are of particular interest for allele specific differences in endocytic assembly and trafficking because, as professional antigen presenting cells (APCs), they have specialized endosomal trafficking mechanisms not possessed by other cells that preserve internalized antigen for assembly with class I molecules before T cell epitopes are destroyed by excessive lysosomal degradation (Accapezzato et al., 2005, Chatterjee et al., 2012, Alloatti et al., 2015). Although some studies have suggested roles for endosomal recycling and assembly in constitutive HLA-I induction (MacAry et al., 2001), the major immunologically relevant role for HLA-I recycling is thought to be in antigen cross-presentation. Antigen cross-presentation is a process by which exogenous antigens derived from soluble proteins or dying cells are internalized by APCs, processed, and presented on HLA-I molecules. The assembly pathways for cross-presentation are highly debated but can be broadly split into two categories: vacuolar and cytosolic. Vacuolar cross-presentation involves endocytosis or phagocytosis of antigen, degradation of the antigen by endo-lysosomal proteases, and trafficking of HLA-I molecules to the antigen-containing endosome for assembly with peptide and recycling to the cell surface (Colbert et al., 2020). The cytosolic pathway, by contrast, involves export of the protein antigen out of the phagosome/endosome by multiple pathways, cytosolic proteasomal degradation, and transport of processed peptides by TAP either into the ER for standard assembly (Colbert et al., 2020), or back into endosomes/phagosomes for assembly with recycling HLA-I molecules (recent studies have demonstrated TAP redistribution to endosomes and phagosomes in dendritic cells under certain conditions) (Colbert et al., 2020). In both pathways, there is potential for endosomal or phagosomal HLA-I peptide assembly, and many organelles implicated in cross-presentation intersect with HLA-I recycling pathways.

HLA-I alleles are linked to favorable and unfavorable outcomes in disease (Carrington and Walker, 2012, Matzaraki et al., 2017). In HIV infections, recent studies point to an association between control of HIV and tapasin-independence, linking unconventional assembly mechanisms with clinical outcomes (Bashirova et al., 2020). While such associations implicate HLA-I assembly characteristics as an important factor underlying immune response differences, the specific mechanisms that contribute to different disease outcomes remain to be further elucidated. It is clear that assembly pathways and the impact of HLA-I polymorphisms vary between different cell types (Yarzabek et al., 2018). Particularly for professional APCs, where unconventional assembly pathways have been studied previously in the context of cross-presentation, it is not understood how HLA-I polymorphisms affect assembly with endogenous or exogenous antigens within endo-lysosomal compartments. While these compartments are well-known sites of HLA class II assembly, they generally lack ER chaperones and factors important for HLA-I assembly, necessitating more flexibility in permissible assembly conditions. These questions were addressed in this study, using primary human monocytes and monocyte-derived dendritic cells (moDCs), and a focus on endo-lysosomal assembly characteristics of B*08:01 and B*35:01. Unlike B*35:01, B*08:01 follows the canonical ER pathway for its assembly and is strongly dependent on tapasin and TAP for cell-surface expression (Rizvi et al., 2014, Geng et al., 2018b, Yarzabek et al., 2018, Bashirova et al., 2020). Comparisons of endo-lysosomal localization and assembly of HLA-B in monocytes and moDCs from donors expressing these two allotypes were undertaken. B*08:01 and B*35:01 were studied as representative members of groups of allotypes that, respectively, favor canonical or non-canonical assembly modes.

## RESULTS

### Low cell-surface HLA-Bw6 half-lives in moDCs and allotype-dependent differences in cell surface stability and peptide occupancy

We addressed the hypothesis that the ability of B*35:01 to assemble independently of PLC components combined with its suboptimal ER assembly (Rizvi et al., 2014, Geng et al., 2018b, Yarzabek et al., 2018, Bashirova et al., 2020) could enable greater endosomal assembly compared to an allotype such as B*08:01 that conforms to the canonical TAP and tapasin-dependent pathway for optimal peptide loading (Rizvi et al., 2014, Geng et al., 2018b, Yarzabek et al., 2018, Bashirova et al., 2020). To address this hypothesis, donors who are B*08:01^+^ or B*35:01^+^ were selected for comparisons (Groups 1 and 2, **Table S1**) from our previously described cohort of HLA genotyped healthy participants (Yarzabek et al., 2018). HLA-B allotypes can be grouped into either the HLA-Bw4 or HLA-Bw6 serotypes, which are recognized by specific antibodies. For the first set of analyses, donors were selected who are heterozygous for HLA-B alleles encoding one HLA-Bw6 and one HLA-Bw4 allotype. Both B*08:01 and B*35:01 contain the Bw6 epitope, which can be detected with an anti-Bw6 monoclonal antibody, the full specificities of which we have previously described (Yarzabek et al., 2018). This antibody does not recognize HLA-A. Some HLA-C allotypes are recognized by anti-Bw6, but donor selection ensured the absence of more than one cross-reactive HLA-C for each HLA-B with a Bw6 epitope (Groups 1 and 2, **Table S1**). Previous studies suggest that HLA-C protein expression in PBMCs is at least 4-6 fold lower than HLA-B (Apps et al., 2015). The significant dominance of HLA-B protein expression relative to HLA-C protein expression (> 4-fold for surface and >3-fold for total) was confirmed by comparisons of anti-HLA-Bw6 signals in moDCs from three sets of donors expressing either HLA-B or HLA-C allotypes with Bw6 epitopes (**Figure S1**). Thus, in the selected donors (Groups 1 and 2, **Table S1**), the dominant signals measured by anti-Bw6 arise from B*08:01 or B*35:01, which are equivalently recognized by anti-Bw6 (Yarzabek et al., 2018).

We determined the cell surface half-lives of HLA-Bw6 on the surface of B*35:01^+^ or B*08:01^+^ moDCs by treating with brefeldin A (BFA) and measuring the surface decay-rates over a four-hour time course. BFA treatment blocks forward trafficking of proteins from the ER to the cell surface by disrupting the Golgi complex (Fujiwara et al., 1988); as a result, this assay measures how rapidly HLA-B is endocytosed from the cell surface (**Figure 1A**). B*08:01^+^ moDCs have a longer Bw6 half-life compared to B*35:01^+^ moDCs, which is in line with previous observations in lymphocyte subsets (Yarzabek et al., 2018). The average half-life for moDC HLA-Bw6 is about two hours, with a shorter B*35:01 half-life compared to B*08:01 (**Figure 1B**). Interestingly, we have previously measured the HLA-Bw6 half-life on monocytes to be about 10 hours (Yarzabek et al., 2018), indicating that moDCs internalize HLA-Bw6 more rapidly than monocytes (**Figure S2**). Thus, in the immature state, dendritic cells appear predisposed to rapid internalization of HLA-I, particularly of B*35:01. Rapid HLA-Bw6 internalization in moDCs compared to other leukocytes could relate to altered dynamics of endocytosis, sorting, and recycling.

**Figure 1.**
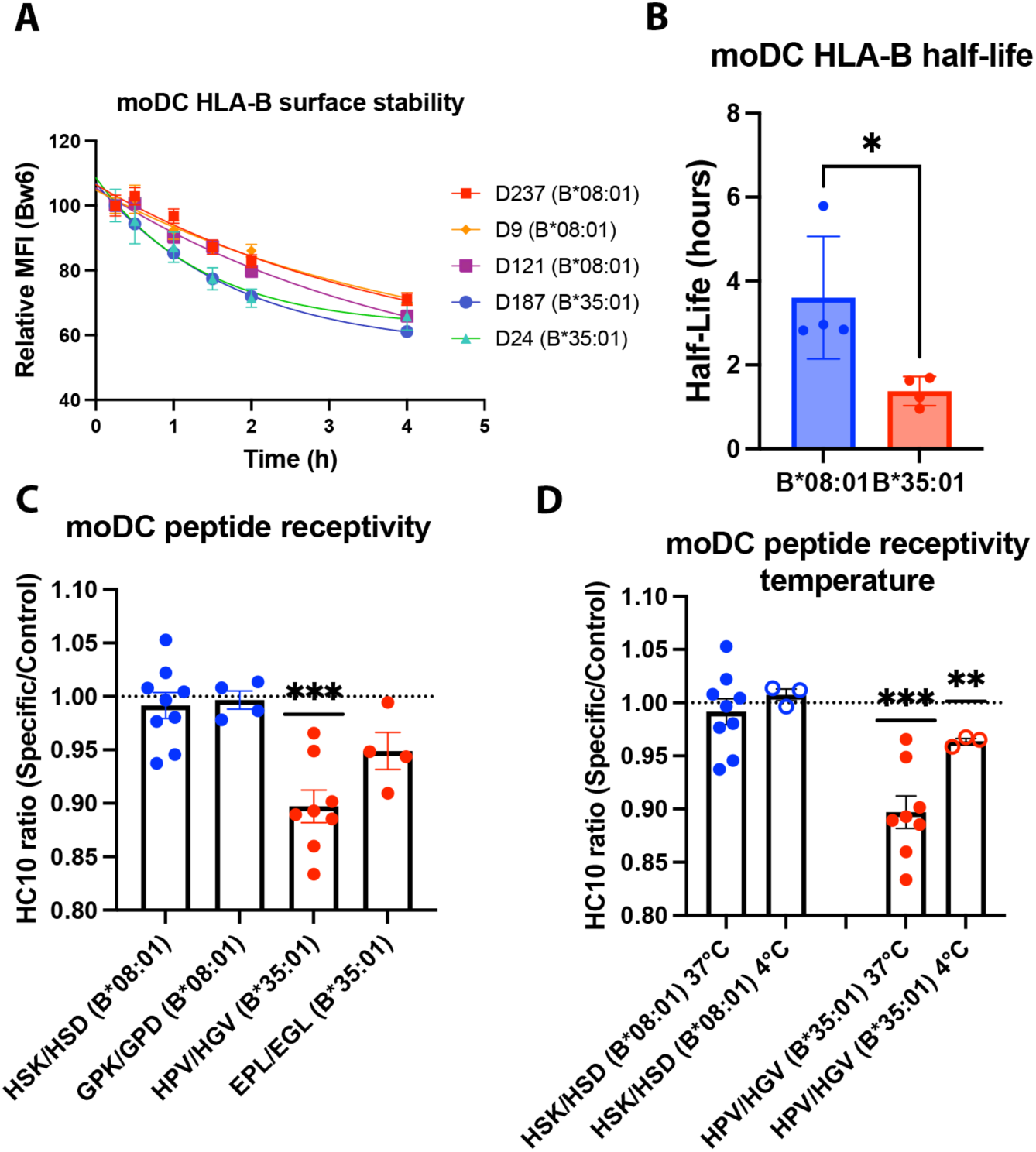
Allotype-dependent differences in HLA-Bw6 surface stability and peptide occupancy in monocyte-derived dendritic cells. **(A)** Representative moDC HLA-Bw6 surface decay over 4-hour time course after treatment with Brefeldin A (BFA). **(B)** Average HLA-Bw6 half-life on moDCs extrapolated from BFA decay-rates. B*35:01^+^ donors were: 24, 187 (n=2), and 210. B*08:01^+^ donors were: 9, 121, 198, and 237. B*08:01^+^ n=4 experiments (4 donors), B*35:01^+^ n=4 experiments (3 donors). Data were analyzed with an unpaired t test. **(C)** moDCs pulsed with B*08:01-specific peptides HSK and GPK or B*35:01-specific peptides HPV and EPL were stained with the monoclonal antibody HC10 to measure peptide-receptive HLA-B. Staining with specific peptides was plotted as a ratio to the control peptides HSD and GPD (for B*08:01) or HGV and EGL (for B*35:01). Ratios were compared to 1 (no difference between specific and control peptides) using a one sample t test. B*08:01 donors were: 94, 121, 130, 148, 178, 198, and 237. B*35:01 donors were: 24, 168, 187, and 210. B*08:01 HSK/HSD n=9, GPK/GPD n=4. B*35:01 HPV/HGV n=8, EPL/EGL n=4. **(D)** moDC peptide receptivity experiments as in (C) including incubation with peptide at 4°C. n=3 experiments at 4°C for each allotype.

Suboptimally assembled MHC-I molecules are subject to cell surface quality control and are more rapidly internalized (Zagorac et al., 2012). To assess whether more rapid loss of cell surface HLA-B in moDCs is related to suboptimal assembly, we employed a peptide occupancy assay previously developed by our lab (Yarzabek et al., 2018), which is based on the higher peptide receptivity of suboptimally loaded MHC-I (Ljunggren et al., 1990, Schumacher et al., 1990). B*08:01 or B*35:01-specific peptides were incubated with moDCs for four hours. Additionally, peptides that were truncated at the C terminus and mutated at key anchor residues were used as controls for non-binding peptides. A monoclonal antibody, HC10, was used to detect empty conformers of HLA-I. A reduction in HC10 signal upon addition of specific peptide relative to control peptide measures allotype-specific peptide binding (Yarzabek et al., 2018). When moDCs were pulsed with two sets of specific and control peptides, B*35:01 was in general receptive to peptide, whereas B*08:01 was not (**Figure 1C**). A similar trend was previously observed in lymphocytes (Yarzabek et al., 2018) that have higher HLA-I half-lives compared with moDCs (**Figure S2**); thus, more rapid internalization of HLA-Bw6 in moDCs compared to other leukocytes is not explained by suboptimal peptide loading *per se*. Additionally, B*35:01 peptide receptivity on moDCs is somewhat temperature sensitive, as incubation with peptide at 4°C reduced the binding (**Figure 1D**). As 4°C incubation inhibits endocytic trafficking, this indicates that at least a portion of the peptide binding may occur in the endosomal compartments; however, B*35:01 is still more peptide receptive at 4°C than B*08:01, indicating that both on the surface and within endosomes, B*35:01 can bind exogenous peptide more efficiently than B*08:01.

### HLA-B is differently distributed within endo-lysosomal compartments of monocytes and moDCs

To further examine endo-lysosomal distributions of HLA-Bw6 in monocytes and moDCs, we conducted confocal microscopy studies of these cells and quantified co-localization with three markers: EEA1, a marker of early endosomes, Rab11, a marker of recycling endosomes and a previously described endosomal storage compartment in DCs (Montealegre and van Endert, 2018), and LAMP1, a lysosomal marker. HLA class I is known to traffic constitutively through the endosomal system, first entering the EEA1^+^ early endosomes and from there either recycling through the Rab11^+^ recycling endosome back to the surface or routing to the lysosome for degradation. Additional trafficking pathways have been described for DCs and other professional antigen-presenting cells, particularly branching from a Rab11^+^ perinuclear storage compartment (reviewed in (Montealegre and van Endert, 2018)). This compartment has been implicated as a source of HLA-I for vacuolar cross-presentation pathways (Nair-Gupta et al., 2014), and thus we hypothesized that Rab11^+^ endosomes would be a major compartment of HLA-B accumulation and endosomal assembly, particularly for B*35:01.

Monocytes and moDCs were plated onto coverslips, fixed, and stained for HLA-Bw6 along with either EEA1, Rab11, or LAMP1. Representative images for monocytes and moDCs are shown in **Figure 2A** and **2B**, respectively. Monocytes are smaller and rounded compared to moDCs, which have more cytoplasm and some spindle-shaped extensions on the edges. Endocytic HLA-Bw6 in monocytes dominantly localizes to the lysosomes, as both B*08:01^+^ and B*35:01^+^ cells have the greatest co-localization with LAMP1 (**Figure 2C** and **2D**). Monocyte HLA-Bw6 is least co-localized with EEA1. In moDCs, the endocytic HLA-Bw6 is more broadly distributed across each compartment, showing the greatest co-localization with Rab11^+^ endosomes (**Figure 2E** and **2F**). Similar co-localization trends were observed when the microscopy data was analyzed by the Pearson’s correlation method. Monocytes have high HLA-Bw6 co-localization with both Rab11 and LAMP1 (**Figure S3A** and **S3B**), whereas moDCs have high HLA-Bw6 co-localization with Rab11 (**Figure S3C** and **S3D**).

**Figure 2.**
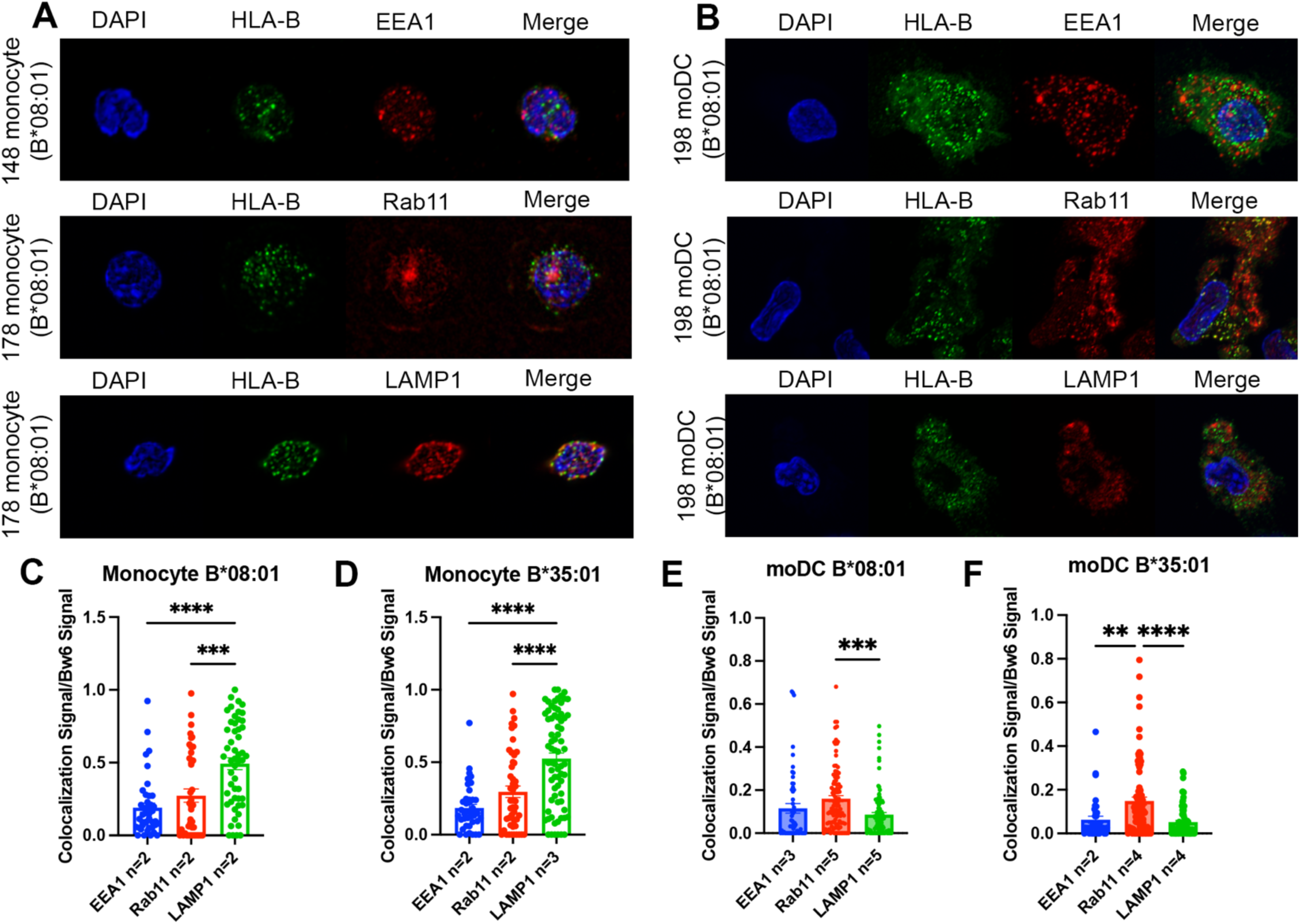
Significant accumulation of HLA-Bw6 in lysosomal (monocytes) or recycling endosomal (moDCs) compartments. **(A)** Representative confocal microscopy images of primary human monocytes stained for HLA-Bw6 co-localization with the early endosome marker EEA1, the recycling endosome marker Rab11, and the lysosomal marker LAMP1. **(B)** Representative moDC staining for HLA-Bw6 co-localization with the same markers. **(C and D)** Monocyte Bw6 co-localization with each indicated marker was quantified by object-based co-localization for 2-4 B*08:01^+^ or B*35:01^+^ donors. **(E and F)** moDC Bw6 co-localization with each of the indicated marker was quantified by object-based co-localization for B*08:01^+^ or B*35:01^+^ donors. Each point is a cell, with at least 20 individual cells imaged per donor and 2-5 donors for each co-localization condition. Co-localization data are represented as the fraction of HLA-Bw6 signal overlapping with the second marker signal. Monocyte B*08:01^+^ donors: 55, 130, 148, and 178. Monocyte B *35:01^+^ donors: 24, 136, 187, and 210. moDC B*08:01^+^ donors: 55, 94, 166, 198, and 237. moDC B*35:01^+^ donors: 24, 168, and 187. One-way ANOVAs were used for analysis to compare the co-localization of HLA-B with each marker.

Together, the results of Figure 2 indicate that monocytes have greater HLA-Bw6 trafficking to lysosomes, indicative of inefficient endosomal assembly and/or more rapid endosomal maturation to lysosomal compartments. Meanwhile, moDCs have most of their HLA-Bw6 localized to Rab11^+^ endosomes, which are an essential storage compartment for HLA-I recruitment to DC antigen^+^ endosomes during cross-presentation (Montealegre and van Endert, 2018). Overall, human moDCs are more poised for HLA-Bw6 endosomal assembly and re-expression on the surface compared to monocytes, which fits the more differentiated and specialized state of moDCs. Cell-specific differences dictate HLA-Bw6 half-lives and intracellular distributions in monocytes and moDCs. Reduced internalization rates (**Figure S2**) and more rapid endosomal maturation in monocytes (**Figure 2C-2D**) could deplete the endosomal HLA-Bw6 pools, whereas more efficient endocytosis (**Figure S2**) but sorting to recycling endosomes (**Figure 2E-2F**) could maintain greater HLA-B endosomal sampling in moDCs.

### moDC B*35:01 is dependent on endo-lysosomal acidification for its surface expression

To better understand the occurrence of HLA-B assembly within endo-lysosomal compartments, monocytes and moDCs were treated over a time course with bafilomycin A1, an inhibitor of the V-ATPase that maintains low endo-lysosomal pH, and surface HLA-B expression was measured by flow cytometry (**Figure 3A, 3B**). Treatment with bafilomycin resulted in an increase in the surface expression of HLA-Bw6 in both B*08:01^+^ and B*35:01^+^ monocytes (**Figure 3C**). These results are in line with the high lysosomal accumulation of HLA-Bw6 observed with confocal microscopy (**Figure 2**); thus, it seems that the increase in lysosomal pH with bafilomycin treatment rescues the HLA-B in this compartment from degradation, particularly in B*08:01^+^ monocytes (**Figure 3C**). Conversely, we observed that HLA-Bw6 expression was unchanged in B*08:01^+^ moDCs after bafilomycin treatment, whereas B*35:01^+^ moDCs decreased their HLA-Bw6 expression (**Figure 3D**). This finding indicates that, unlike in monocytes, low endo-lysosomal pH is important for maintaining B*35:01 surface expression in moDCs. A gradient of V-ATPase subunits is present along the endosomal pathway, with the highest number of active complexes present in the lysosomes and lowest amount present in early endosomes (Lafourcade et al., 2008). Thus, it is likely that assembly and trafficking through the entire endo-lysosomal pathway is perturbed by bafilomycin treatment, which negatively affects B*35:01 in moDCs. In contrast, there is reduced proteasomal dependence of HLA-Bw6 expression in B*35:01^+^ monocytes and moDCs compared with the corresponding B*08:01^+^ cells (**Figure S4A** and **S4B**). These finding suggests that, in line with previously observed tapasin and TAP-independent expression (Rizvi et al., 2014, Geng et al., 2018b), B*35:01 compared with B*08:01 is less reliant upon canonical proteasome-dependent ER assembly, but more reliant on endosomal assembly

**Figure 3.**
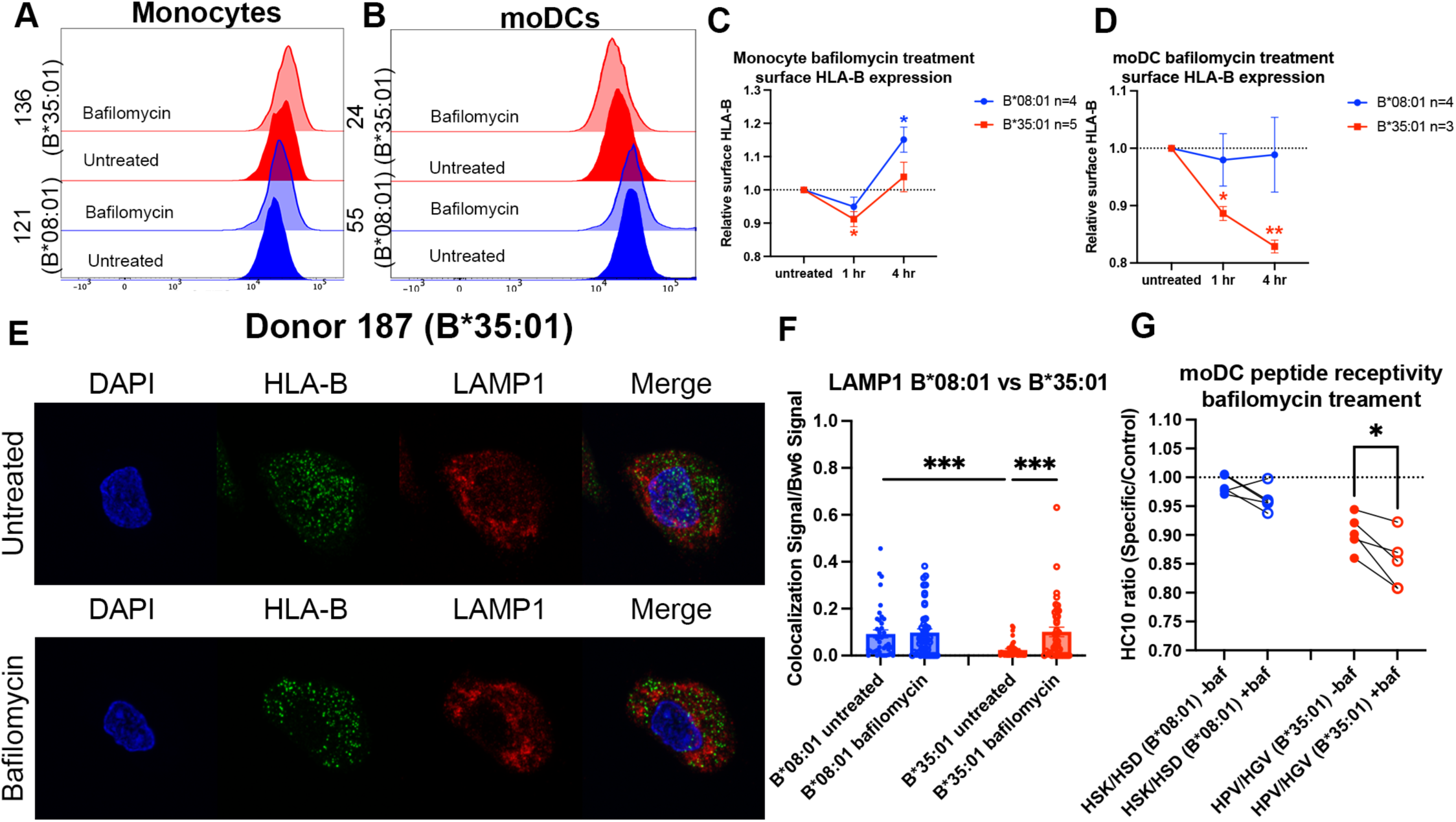
Disruptions to endo-lysosomal pH alter HLA-B*35:01 surface expression and induce lysosomal accumulation in moDCs. Monocytes or monocyte-derived dendritic cells were treated with 200 nM bafilomycin A1 for 1, 2, or 4 hrs. Treatment with bafilomycin was followed by staining for surface markers and HLA-Bw6, followed by analysis via flow cytometry. **(A and B)** Representative HLA-B expression flow cytometry histograms are shown for monocytes **(A)** and moDCs **(B). (C)** Relative changes in HLA-Bw6 expression on the surface of HLA-B*08:01^+^ (n=4) or HLA-B*35:01^+^ (n=5) monocytes over the 4-hour bafilomycin time course. B*08:01 and B*35:01 expression at each time point was compared to the normalized untreated expression with one sample t tests. B*08:01^+^ donors for these experiments: 121, 130 (n=2), and 237. B*35:01^+^ donors for these experiments: 24, 141 (n=2), and 210 (n=2). **(D)** Relative changes in HLA-Bw6 expression on the surface of B*08:01^+^ (n=4) and B*35:01^+^ (n=3) moDCs. B*08:01 and B*35:01 expression at each time point was compared to the normalized untreated expression with one sample t tests. B*08:01^+^ donors for these experiments: 55 (n=2), 166, and 198. B*35:01^+^ donors for these experiments: 24 (n=2) and 187. **(E)** moDC confocal microscopy experiments comparing HLA-Bw6 co-localization with LAMP1 with and without bafilomycin treatment. **(F)** Object-based co-localization quantification of HLA-Bw6 with LAMP1 with and without bafilomycin treatment. B*08:01^+^ donors were 94 and 237, and the B*35:01^+^ donors were 168 and 187 (n=2 for each group). Unpaired t tests were used to compare co-localization with and without bafilomycin. **(G)** Peptide receptivity of B*08:01 and B*35:01 carried out in the presence or absence of bafilomycin. Receptivity of each allotype was compared +/-baf treatment via paired t tests. B*08:01^+^ donors: 94 (n=2), 178, 198, and 237; n=5 independent experiments. B*35:01^+^ donors: 24, 168, 187, and 210 (n=2); n=5 independent experiments.

Next, we treated moDCs with bafilomycin as before, but performed confocal microscopy to examine endo-lysosomal HLA-B redistribution after four hours of bafilomycin treatment. Representative images do not show obvious changes in HLA-Bw6 or LAMP1 distribution throughout the cell after treatment (**Figure 3E**), but quantifications indicate increased HLA-Bw6 co-localization with LAMP1^+^ lysosomes upon treatment with bafilomycin in B*35:01^+^ moDCs, but not in B*08:01^+^ moDCs (**Figure 3F**). Additionally, in the untreated condition, there is higher steady-state HLA-Bw6 localization to lysosomes in B*08:01^+^ moDCs compared with B*35:01^+^ moDCs. Bafilomycin treatment increases HLA-Bw6/LAMP1 co-localization in B*35:01^+^ moDCs to the B*08:01^+^ moDC levels. Bafilomycin treatment generally has no effect on HLA-Bw6/Rab11 co-localization in B*35:01^+^ moDCs (**Figure S4C**), and while object-based co-localization analyses indicate increased co-localization of HLA-Bw6 with Rab11 in B*08:01^+^ moDCs (**Figure S4D**), the Pearson’s correlation analyses do not support this conclusion (**Figure S4E**). Pearson’s correlation analyses confirm more significant increases in B*35:01 localization to lysosomes following bafilomycin treatment (**Figure S4F**).

Finally, bafilomycin treatment further enhances the peptide receptivity of B*35:01 in moDCs, indicating that bafilomycin reduces the normal level of antigen supply or alters assembly efficiency for B*35:01, resulting in more abundant suboptimal conformers (**Figure 3G**). Together, the data of Figure 3 indicate that raising the endo-lysosomal pH via bafilomycin treatment results in reduced surface expression and enhanced lysosomal accumulation of B*35:01, which is coincident with accumulation of more peptide-receptive B*35:01 conformers. HLA-Bw6 surface expression is unaffected by bafilomycin in B*08:01^+^ moDCs. Thus, endosomal pH-dependent processes are required by B*35:01 but not B*08:01 for maintaining optimal constitutive expression in moDCs.

### Monocytes and moDCs differ in exogenous antigen uptake efficiencies and processing pathways

Human moDCs are shown to process and assemble antigens for cross-presentation in their endo-lysosomal (vacuolar) compartments (Tang-Huau et al., 2018), a pathway we predicted would be more permissive for B*35:01 based on the results described in Figures 1-4. Although not much work has been done to investigate antigen processing by monocytes, it is generally believed that differentiation into DCs is required for induction of monocyte cross-presentation function (Doring et al., 2019). Consistent with this view, monocytes and moDCs uptake soluble antigen to different extents, with greater moDC endocytosis of fluorescently labeled bovine serum albumin over a 60-minute time course (**Figure 4A**). To study processing of endocytosed antigen, we used DQ-Ova as a model for antigen degradation. DQ-Ova is a soluble ovalbumin protein labeled with excess, self-quenching BODIPY fluorophores, which become fluorescent upon proteolytic cleavage of the protein. After DQ-Ova pulsing at either 37°C or 4°C for 30 min, the cells were chased in media or media + inhibitors, followed by fixation and flow cytometric analysis. When monocytes were chased with lysosome or proteasome inhibitors and normalized to untreated cells, antigen degradation was completely inhibited by bafilomycin treatment (**Figure 4B**). MG132 inhibition of the proteasome had no effect, while the inhibitor combination inhibited similarly to bafilomycin alone. Thus, monocytes mainly use their lysosomes for exogenous protein degradation.

**Figure 4.**
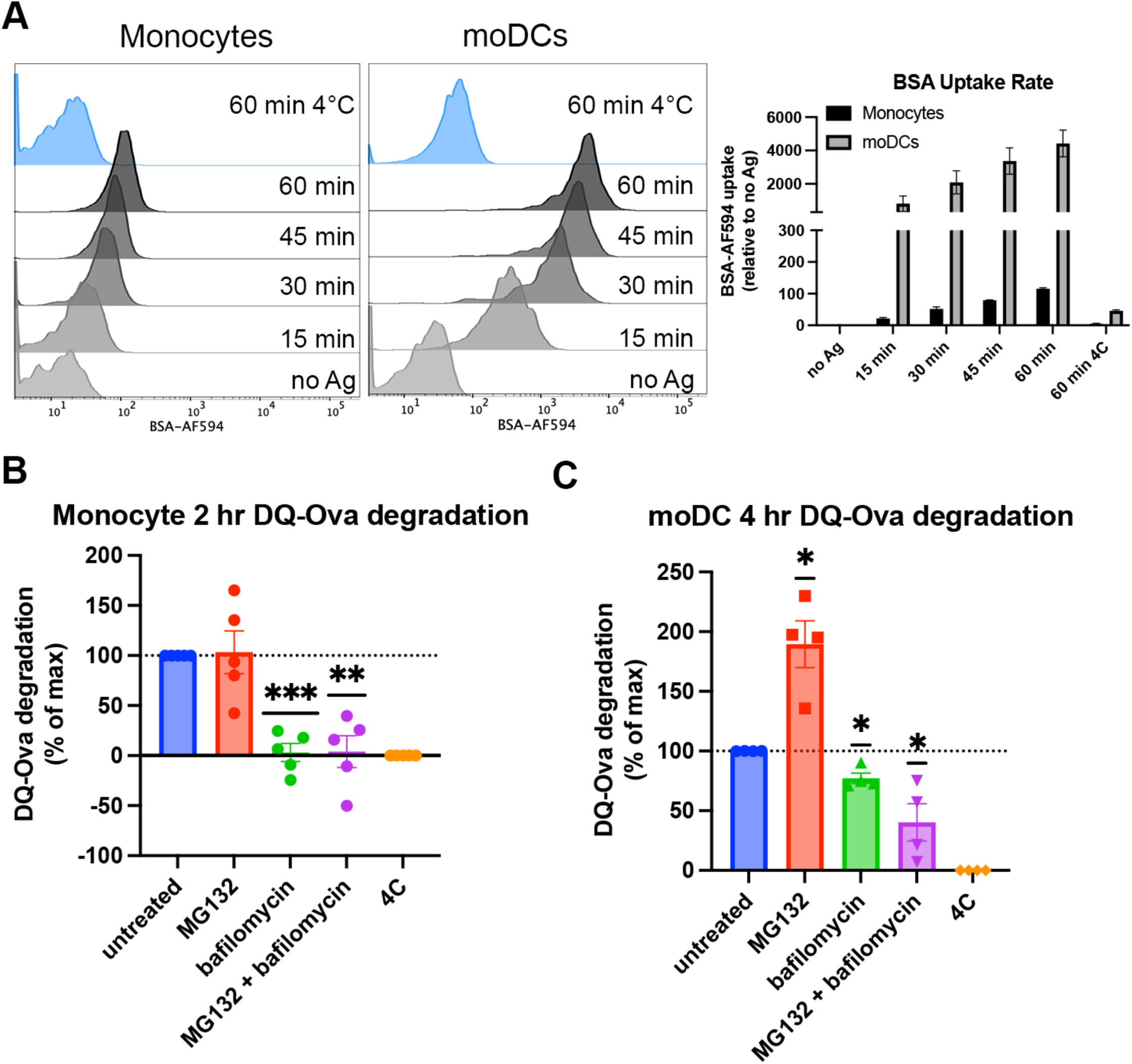
Cell type-dependent differences in antigen uptake and processing pathways. **(A)** Monocytes or moDCs were pulsed with BSA labeled with Alexa fluor 594 for 15-minute intervals, followed by washing, fixation, and flow cytometric analyses of uptake. Representative histogram plots are shown, as well as averaged uptake rates. N=2 independent experiments for each cell type. Monocyte donors: PCD22F, PCD25F. moDC donors: 255, PCD37M. **(B and C)** Assessments of antigen degradation pathways in monocytes **(B)** and moDCs **(C)** were performed using the soluble antigen DQ-Ova. Monocyte DQ-Ova degradation for 2 hrs with inhibitors relative to untreated was quantified in **(B)**, and moDC degradation for 4 hrs with inhibitors was quantified in **(C)**. For monocytes, the experiment was repeated n=5 times, and n=4 times for moDCs. Monocyte donors: 248, 250, 255, 270, and 273. moDC donors: 250, 253, 270, and 275. Data was normalized by subtracting degradation at 4°C from all other conditions and setting degradation at 37°C as the maximum. The effect of each inhibitor on degradation compared to untreated 37°C was assessed with a one sample t test.

In moDCs, MG132 inhibition actually increases the degradation of DQ-Ova, likely due to a compensatory enhancement of lysosome-mediated degradation (Pandey et al., 2007). In contrast to monocytes, bafilomycin did not completely block DQ-Ova degradation, and the combination of MG132 and bafilomycin further reduced the degradation (**Figure 4C**). As bafilomycin increases the lysosomal pH and inhibits most pH-sensitive proteases present in this compartment, the upregulation of lysosomal degradation by MG132 treatment cannot increase the antigen degradation in the MG132 + bafilomycin combination treatment as seen with MG132 alone. Thus, for soluble protein degradation, moDCs use both cytosolic and lysosomal pathways. Additionally, the extent of lysosomal degradation of antigen differs between moDCs and monocytes, as bafilomycin alone inhibits monocyte antigen degradation to a greater extent than moDCs. Increased uptake and reduced proteolysis within the endo-lysosomal compartments upon moDC differentiation could explain why some endocytosed antigen might undergo proteasomal processing in these cells, as more protein may be preserved for export from endosome to cytosol.

### Cross-presentation via B*35:01 is more efficient than B*08:01 even when matched for T cell response sensitivity and is more affected by cathepsin inhibition

Since endo-lysosomal antigen degradation occurs in both monocytes and moDCs, we further examined the model that B*35:01 would have cross-presentation advantages in both monocytes and moDCs due to its increased propensity both for constitutive endo-lysosomal assembly (**Figure 3**) and assembly with exogenous peptides **(Figure 1**). To perform cross-presentation assays with human cells, we took advantage of the fact that memory cytotoxic T lymphocytes (CTLs) specific for Epstein-Barr Virus (EBV) antigens are broadly prevalent in humans. Two EBV proteins contain known epitopes for both B*08:01 (RAKFKQLL (RAK) and FLRGRAYGL (FLR)) and B*35:01 (EPLPQGQLTAY (EPL) and YPLHEQHGM (YPL)). The FLR and YPL epitopes are derived from the EBNA3A protein, and the RAK and EPL epitopes are derived from the BZLF1 protein (Thomson et al., 1995, Rist et al., 2015). We sorted and expanded antigen-specific CTLs with B*08:01-RAK, B*08:01-FLR, B*35:01-EPL, and B*35:01-YPL tetramers from donors expressing the relevant HLA-B allotypes (Group 3, **Table S1**). Peptide titration experiments with the CTLs demonstrated varied sensitivities of each CTL line to peptide, with B*35:01-YPL eliciting the most sensitive response and B*08:01-RAK the least sensitive (**Figure 5A**). B*08:01-FLR and B*35:01-EPL CTLs displayed similar sensitivities to peptide.

**Figure 5.**
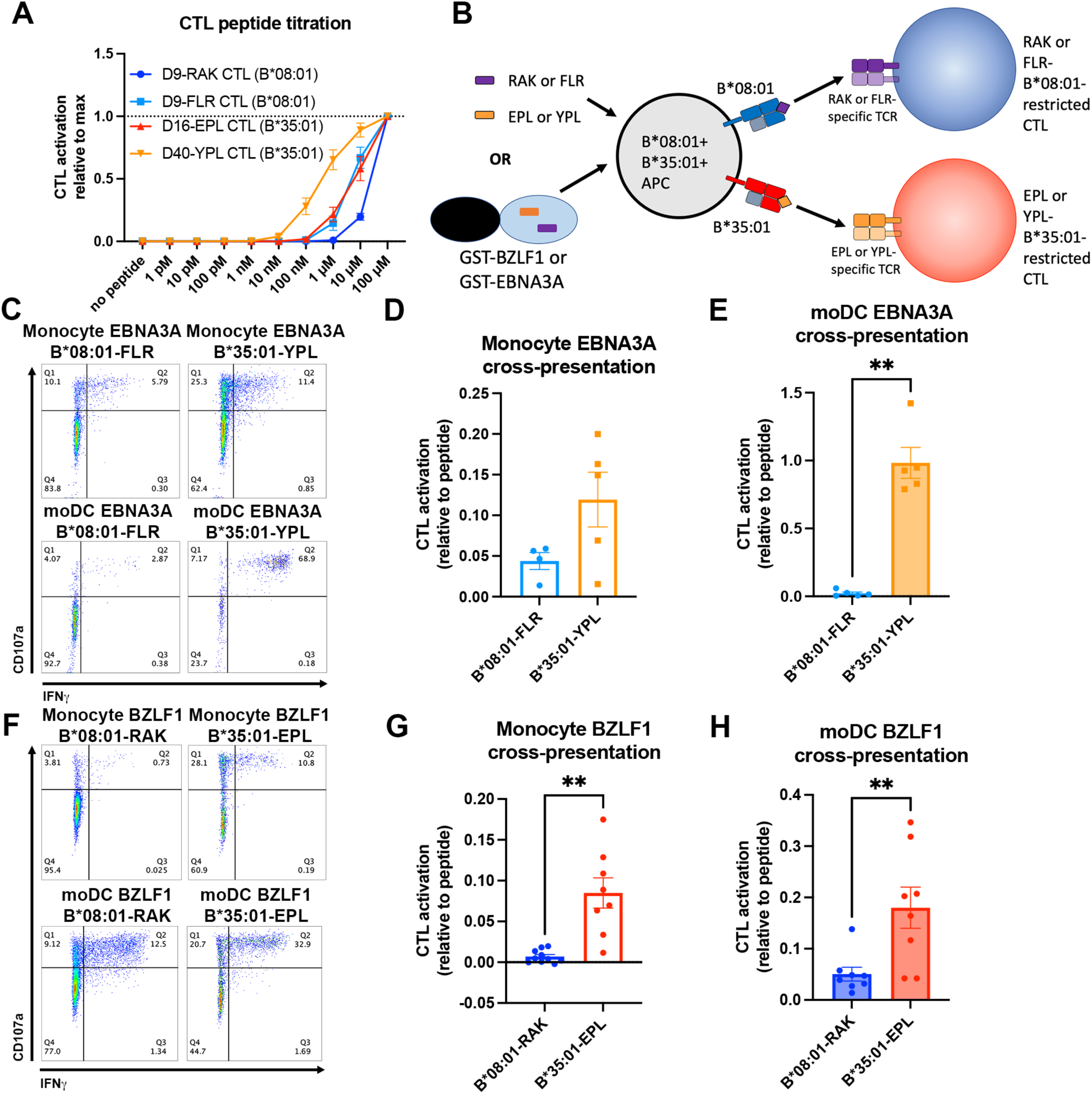
Cross-presentation of epitopes derived from EBV proteins by B*35:01 compared to B*08:01. **(A)** B*08:01-RAK CTLs from donor 9, B*08:01-FLR CTLs from donor 9, B*35:01-EPL CTLs from donor 16, and B*35:01-YPL CTLs from donor 40 were used in peptide titration experiments to measure sensitivity to peptide. PBMCs from B*08:01^+^ or B*35:01^+^ donors were pulsed with peptide overnight at different concentrations, followed by co-culture with each CTL and flow cytometric assessment of activation. B*08:01^+^ PBMC donors: 94 (n=2) and 148 (n=2). B*35:01^+^ PBMC donors: 24 (n=4). N=4 independent experiments for B*08:01-RAK and B*35:01-EPL CTLs, and n=3 experiments for B*08:01-FLR and B*35:01-YPL CTLs. **(B)** Schematic representation of cross-presentation assay. B*08:01^+^/B*35:01^+^ double-positive monocytes or moDCs were pulsed with either 50 µM canonical B*08:01 peptide (RAK or FLR), 50 µM canonical B*35:01 peptide (EPL or YPL), or purified GST-BZLF1 or GST-EBNA3A protein antigen (100 µg) for 6 hrs, then co-cultured with previously expanded CTLs (either B*08:01-restricted or B*35:01-restricted) at a 1:1 CTL:APC ratio for 5 hrs. CTLs were assessed for activation by surface CD107a expression and intracellular IFNγ. **(C)** Representative flow cytometry plots are shown for CTLs co-cultured with monocytes and moDCs during EBNA3A cross-presentation. CD107a degranulation and intracellular IFNγ expression were measured by flow cytometry. **(D)** Monocyte cross-presentation of EBNA3A quantified as a ratio relative to peptide, n=4 B*08:01-FLR experiments, n=5 B*35:01-YPL experiments. **(E)** moDC cross-presentation of EBNA3A quantified as a ratio relative to peptide, n=5 experiments. **(F)** Representative flow cytometry plots are shown for CTLs co-cultured with monocytes and moDCs during EBNA3A cross-presentation. **(G)** Monocyte cross-presentation of BZLF1 quantified as a ratio relative to peptide, n=9 experiments. **(H)** moDC cross-presentation of BZLF1 quantified as a ratio relative to peptide, n=7 experiments. B*08:01 and B*35:01 cross-presentation in D, E, G, and H compared with paired t tests. Monocyte and moDC APC donors were: 16, 25, and 132. B*08:01-RAK CTL donors: 9 and 16. B*08:01-FLR CTL donor: 9. B*35:01-EPL CTL donor: 16. B*35:01-YPL CTL donor: 40.

To control for donor-to-donor antigen uptake and processing differences during cross-presentation assays, monocytes and moDCs were used from donors expressing both B*08:01 and B*35:01 so that antigen presentation via both allotypes occurs within the same cells (Group 4, **Table S1**). The FLR, RAK, YPL, and EPL peptides were used as positive controls for CTL activation, and the purified recombinant EBNA3A or BZLF1 proteins were used as model soluble antigens for cross-presentation (**Figure 5B**). CTL activation was measured by surface CD107a expression (a marker of CTL degranulation) and intracellular IFNγ production.

As noted above, resting CD14^+^ monocytes are generally considered to not be competent for cross-presentation (Doring et al., 2019), instead requiring differentiation to moDCs. Surprisingly, however, there is a readily detectable CTL activation in response to EBNA3A cross-presentation, for both the B*08:01-FLR and B*35:01-YPL epitopes (**Figure 5C**). In monocytes, despite the higher percentage of activated B*08:01-FLR CTL activation in response to peptide (**Figure S5A**), B*35:01-YPL CTLs were activated to a greater extent with EBNA3A (**Figure S5B**). Normalization of EBNA3A-induced activation to peptide-induced activation confirmed the trend of greater B*35:01-YPL cross-presentation efficiency (**Figure 5D**). These patterns were exaggerated for moDC presentation of peptide and EBNA3A antigen (**Figure S5C** and **S5D**), where B*35:01-YPL cross-presentation was greatly enhanced compared to monocytes (**Figure 5C**), approaching the level of peptide-induced activation (**Figure 5E**). Thus, in both monocytes and moDCs, the EBNA3A YPL epitope is cross-presented via B*35:01 more efficiently than the FLR epitope via B*08:01.

As the B*35:01-YPL response was sensitive to very low doses of peptide (**Figure 5A**), we sought to confirm our cross-presentation findings with the BZLF1 antigen, which contains RAK, the B*08:01 epitope, and EPL, the B*35:01 epitope. Peptide activation of both BZLF1 epitopes is less sensitive compared to their EBNA3A counterparts (**Figure 5A**), offering conditions to examine cross-presentation where antigen may be more limiting. Cross-presentation of the RAK epitope was minimal in monocytes but more detectable in moDCs, while the EPL epitope was more readily cross-presented in both cell types (**Figure 5F**). While there are no allotype-dependent differences in peptide-mediated activation in monocytes (**Figure S5E**), the B*35:01-EPL epitope from the whole BZLF1 antigen activates CTLs to a greater extent than the B*08:01-RAK epitope within the same experiments and same donor APC (**Figure S5F**). The B*35:01-EPL advantage for cross-presentation persists when normalized to peptide activation levels within each experiment (**Figure 5G**). In moDCs, there is greater activation of B*35:01-restricted CTLs with both peptide and BZLF1 compared to B*08:01 (**Figure S5G** and **S5H**), and when BZLF1 activation is normalized to peptide activation the advantage of B*35:01-EPL CTLs over B*08:01-RAK CTLs for cross-presentation persists (**Figure 5H**).

In comparing the relative cross-presentation efficiencies of all four epitopes in monocytes and moDCs, it is notable that the epitope with the lowest response sensitivity (B*08:01-RAK, **Figure 5A**) displays the lowest cross-presentation efficiency in monocytes, whereas the epitope with the highest response sensitivity (B*35:01-YPL, **Figure 5A**), displays the highest cross-presentation efficiency in moDCs (**Figures 6A and 6B**). Notably, however, in comparing epitopes with similar response sensitivities (B*08:01-FLR vs B*35:01-EPL; **Figure 6A**), the B*35:01 cross-presentation advantage over B*35:01 is still apparent, particularly in moDCs (**Figure 6C**).

**Figure 6.**
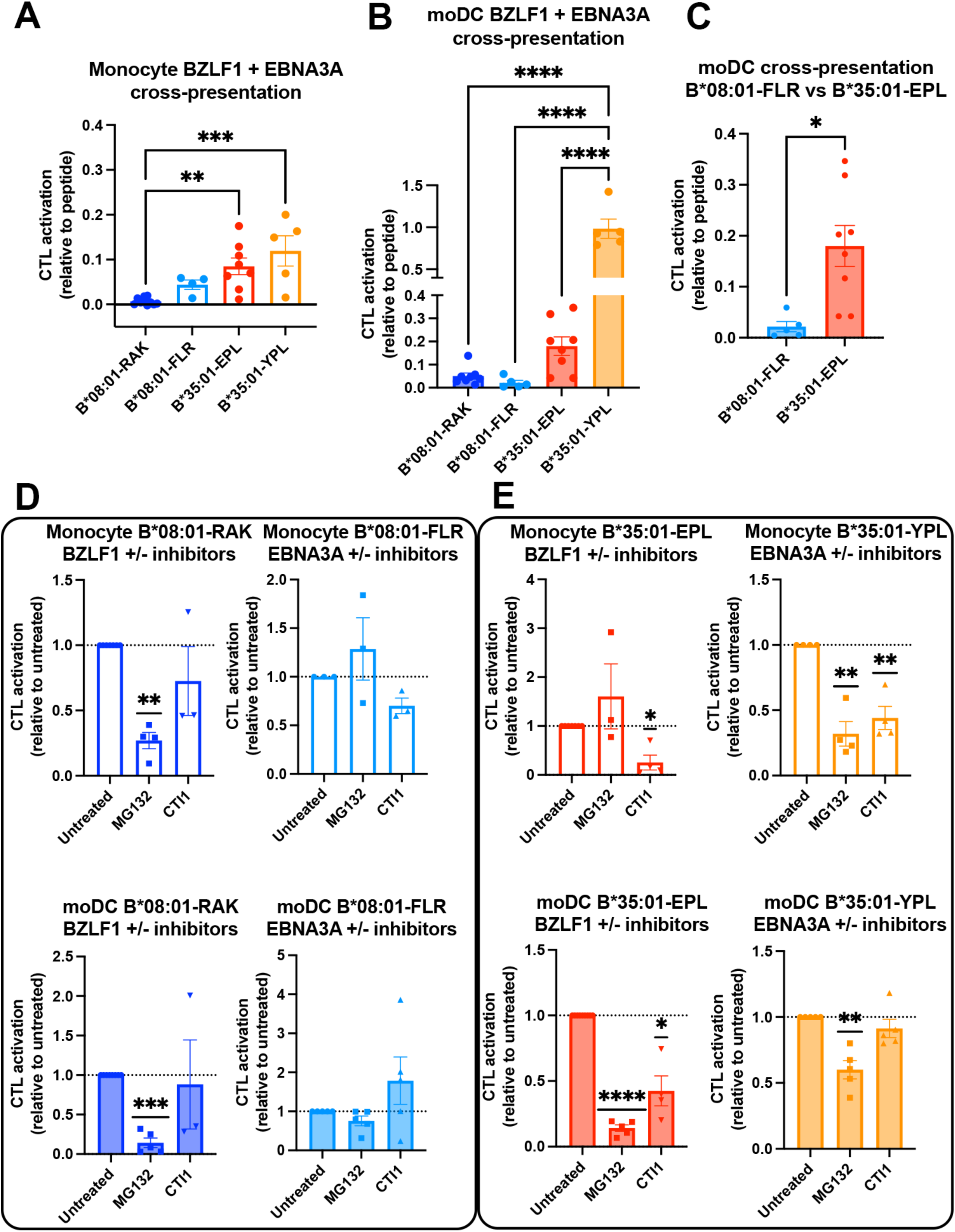
Cross-presentation by B*35:01 displays greater sensitivity to cathepsin inhibition and is more efficient than B*08:01 when matched for T cell responsiveness. **(A and B)** Cross-presentation efficiencies of B*08:01-RAK, B*08:01-FLR, B*35:01-EPL, and B*35:01-YPL epitopes from BZLF1 or EBNA3A were compared in monocytes **(A)** or moDCs **(B)**. Differences were assessed by One-Way ANOVA analysis. **(C)** Two epitopes with similar CTL sensitivities to peptide **(Figure 5A)** were compared for cross-presentation efficiencies in moDCs with an unpaired t test, displaying a trend towards more efficient B*35:01-EPL cross-presentation. **(D-E)** Cross-presentation assays were performed as previously described, with the addition of inhibitor treatment. During monocyte or moDC pulse with 100 µg protein antigen, either MG132 or Cathepsin Inhibitor 1 (CTI1) were added to the APCs to inhibit different pathways of antigen processing. Cross-presentation with inhibitors was compared to untreated with a one-sample t test. For monocyte cross-presentation: n=4 MG132 and n=3 CTI1 B*08:01-RAK treatments, n=3 MG132 and CTI1 B*08:01-FLR treatments, n=3 MG132 and n=4 CTI1 B*35:01-EPL treatments, and n=4 MG132 and CTI1 B*35:01-YPL treatments. For moDC cross-presentation: n=5 MG132 and n=3 CTI1 B*08:01-RAK treatments, n=5 MG132 and CTI1 B*08:01-FLR treatments, n=5 MG132 and n=4 CTI1 B*35:01-EPL treatments, and n=5 MG132 and CTI1 B*35:01-YPL treatments.

We further tested the sensitivities of each CTL response to MG132 inhibition of the proteasome, which probes the relevance of the cytosolic pathway of antigen processing and presentation, and to treatment with the Cathepsin Inhibitor I, an inhibitor of cathepsins B, K, L, and S (**Figure 6D** (B*08:01) and **6E** (B*35:01)). With the exception of the B*08:01-FLR epitope in monocytes, cross-presentation by B*08:01 generally displayed sensitivity to proteasome inhibition even including in monocytes for the B*08:01-RAK epitope (**Figure 6D**). In contrast, B*35:01 responses displayed significant sensitivity to either the Cathepsin Inhibitor I alone or to both MG132 and the Cathepsin Inhibitor I (**Figure 6E**). Although significance was not achieved for cathepsin inhibition of the B*35:01-YPL response in moDCs, there was a trend towards inhibition. The generally greater sensitivity to CTI1 treatment indicates that B*35:01 uses the vacuolar (endo-lysosomal) pathway of cross-presentation exclusively or in combination with the cytosolic pathway depending on cell type. The ability to use both the vacuolar and cytosolic pathways of assembly for generating B*35:01-EPL epitopes could explain the higher efficiency of B*35:01-EPL cross-presentation relative to B*08:01-FLR in moDCs.

## DISCUSSION

Our findings, along with previous studies on HLA class I polymorphisms, provide evidence that assembly characteristics associated with various allotypes confer unique advantages and disadvantages for antigen presentation, depending on the cellular environment. In our study, we observe that human monocytes and moDCs differ in their endo-lysosomal pathways, and that within both cell types, B*08:01 and B*35:01 vary in their utilization of these pathways (**Figure 7**). First, HLA-B is rapidly internalized from the surface of moDCs, with a shorter average cell surface half-life for B*35:01 compared with B*08:01 (**Figure 1**). The short HLA-B half-lives in moDCs are likely explained by altered endocytic, sorting and recycling dynamics, as the surface HLA-B half-life is much lower in moDCs than monocytes (**Figure S2)**. Furthermore, moDC B*35:01 is peptide-receptive (**Figure 1**) while B*08:01 is not, indicating that suboptimal peptide loading of HLA-I is not a global characteristic of moDCs that accounts for shorter moDC half-lives. While previous studies have shown that optimal peptide loading and complex conformation are one set of determinants for cell surface HLA-I stability (Ljunggren et al., 1990, Schumacher et al., 1990), the present findings indicate that internalization and recycling dynamics, which are cell-type dependent, are also key determinants of cell surface HLA-I residence time. These studies also place into context our previous findings that relative surface expression levels of individual HLA-B molecules are both cell and allotype dependent (Yarzabek et al., 2018), a complex product of individual allotype-dependent assembly characteristics and cellular features. Thus, all relative HLA-I expression measurements must be defined in the context of specific cell types.

**Figure 7.**
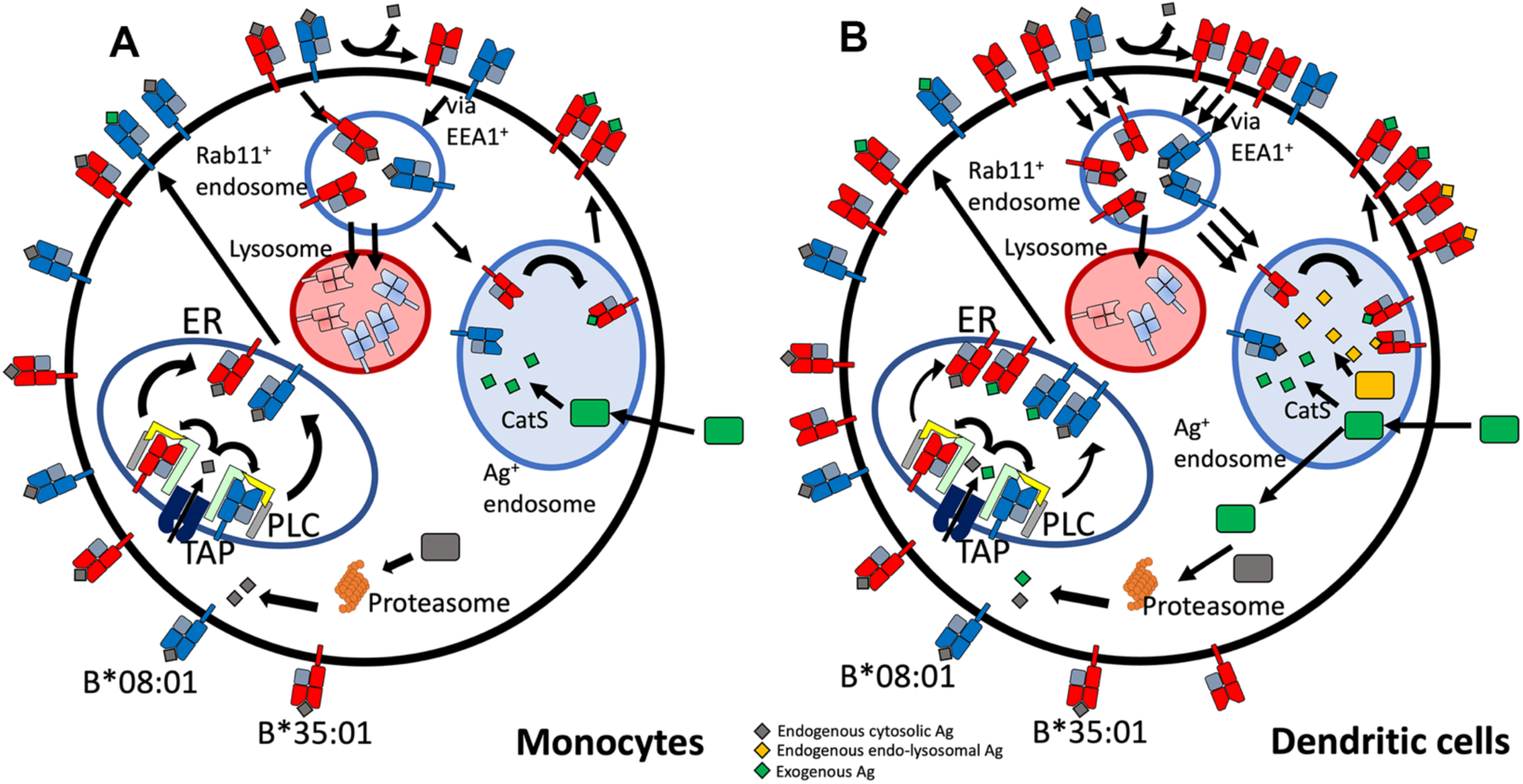
Model of monocyte and moDC endo-lysosomal HLA-B assembly pathways. Monocytes and moDCs have varying HLA-B endo-lysosomal HLA-I trafficking and assembly pathways. **(A)** Endocytosed HLA-Bw6 in monocytes is rapidly trafficked to lysosomes for degradation. Blocking degradation via bafilomycin rescues surface expression. Monocytes can cross-present antigen via a vacuolar pathway in a cathepsin S-dependent manner, which is more permissive for B*35:01, whereas the cytosolic pathway, involving antigen translocation from the endosomes to cytosol, is inefficient in monocytes. **(B)** moDC rapidly internalize HLA-Bw6 through more active endocytic processes. moDCs accumulate HLA-Bw6 in Rab11^+^ endosomes. B*35:01 also displays lower cell surface stability, and requires endo-lysosomal processing for maintenance of its constitutive surface expression, indicating assembly with endogenous endo-lysosomal antigens. moDCs generally cross-present at a higher efficiency than monocytes and can utilize both the cytosolic and vacuolar pathways of antigen processing. B*08:01 can cross-present at a higher efficiency in moDCs compared to monocytes, but B*35:01 maintains its advantage over B*08:01 for cross-presentation efficiency, even when matched for T cell response sensitivity. This is likely because of its ability to efficiently assemble via both the cytosolic and vacuolar pathways.

While it is unclear what specialized factors contribute to the enhanced endocytosis of HLA-Bw6 in moDCs, this is likely a mechanism to promote endo-lysosomal antigen sampling. This hypothesis is validated by the greater co-localization of HLA-Bw6 with Rab11^+^ endosomes in moDCs compared to other endo-lysosomal markers (**Figure 2**). This differs from monocytes, which have the most HLA-Bw6 co-localization with LAMP1^+^ lysosomes. Greater lysosomal HLA-B localization in monocytes may be a result of enhanced endosome maturation to lysosomes and degradation of endocytosed conformers, as evidenced by the increase in surface HLA-Bw6 upon bafilomycin treatment in Figure 3. Monocytes thus appear more poised for lysosomal degradation due to more rapid endosome to lysosome maturation, which is disrupted by bafilomycin treatment (Bayer et al., 1998).

In contrast to monocytes, moDCs appear more poised for HLA-B endosomal assembly, particularly for allotypes such as B*35:01. As Rab11^+^ endosomes are important storage compartments for MHC-I assembly in APCs (Montealegre and van Endert, 2018), the pool of HLA-B present here is indicative of a change in the endosomal system from monocytes for more efficient assembly. Indeed, our data suggest that bafilomycin treatment decreases B*35:01 surface expression because of an inhibitory effect on assembly, as the treatment results both in increased peptide-receptive B*35:01 complexes and greater lysosomal B*35:01 accumulation (**Figure 3**). These findings are the first to our knowledge to demonstrate allotype-dependent differences in constitutive endo-lysosomal assembly of bulk HLA-I proteins (**Figure 7**). Previous studies have demonstrated MHC-I assembly with endogenous transmembrane proteins (Tiwari et al., 2007) and endogenous HSV-1 antigens (English et al., 2009) via the endo-lysosomes, and we predict in the context of our findings that presentation of these and other related antigens is allotype-dependent.

Beyond elucidating differences in endogenous HLA-B assembly in the endo-lysosomal system, our findings extend to assembly differences with exogenous antigen (**Figures 5** and **6**). The efficiency of cross-presentation is generally higher for B*35:01 when comparing responses to the same exogenous antigens in the same cells (**Figures 5C-5H**), and when comparing across antigens matched for response sensitivities (**Figure 6C**). Furthermore, monocyte B*35:01-EPL, B*35:01-YPL and moDC B*35:01-EPL cross-presentation responses all display significant sensitivity to cathepsin inhibition (**Figure 6E**), whereas B*08:01 responses are generally more sensitive to MG132 inhibition (with the exception of the monocyte B*08:01-FLR response) (**Figure 6D**). These results suggest that B*35:01 compared to B*08:01 can better exploit vacuolar degradation in monocytes to acquire exogenous antigen, and thus has greater ability to use multiple (cytosolic and vacuolar) cross-presentation pathways. The locations of processing and assembly may be matched as well; while cathepsins are generally found in lysosomes, cathepsin S has been found located in early endosomal compartments of monocytes and macrophages (Schmid et al., 2002). The overall characteristics of B*35:01 that confer the advantages for endo-lysosomal assembly with both endogenous and exogenous antigens include suboptimal ER assembly, resulting in lower cell surface stability, and increased endocytosis followed by binding of peptide-receptive forms of B*35:01 to endosomally-localized antigens (**Figure 7**). Endosomal assembly may also be facilitated by the capability of HLA-B*35:01 to assemble independently of chaperones and factors that are primarily ER localized.

Aside from allotype-dependent differences in cross-presentation efficiencies and pathways, our studies provide important insights into cross-presentation differences between monocytes and moDCs. Whereas resting monocytes are considered non-permissive for cross-presentation (Doring et al., 2019), we show their capability for cross-presentation just a few hours after exposure to antigens purified from *E. coli*. Furthermore, the occurrence of more extensive endo-lysosomal antigen degradation in monocytes (**Figure 4**) favors cross-presentation of epitopes with higher response sensitivities—those that elicit responses at lower antigen doses (**Figures 5A and 6A**). On the other hand, moDCs are more efficient than monocytes at cross-presentation for most antigens, and antigens with high response sensitivities appear to be better able to exploit the specialized DC environment to achieve high cross-presentation efficiency (**Figures 5A and 6B**). Both monocytes and moDCs are, to different extents, permissive for both the vacuolar and cytosolic cross-presentation pathways, depending on epitopes.

Altogether, based on studies of individual HLA-I allotypes in primary human cells, our findings provide evidence that HLA-I polymorphisms determine not only the specificities of peptide presentation and antigen receptor binding (both innate and adaptive (Djaoud and Parham, 2020)), but also influence antigen sampling in specific subcellular compartments (**Figure 7**). We suggest that certain HLA-I allotypes are predisposed for bulk constitutive assembly within endo-lysosomes (**Figure 3**) following suboptimal assembly in the ER (**Figure 1**), and possibly based on their enhanced capability to assemble via non-canonical, ER assembly factor-independent pathways. Thus, the subcellular localization of assembly for some HLA-I allotypes in part overlaps with the sites for HLA class II assembly (Blum et al., 2013). Whereas in the HLA class II pathway, the invariant chain-derived CLIP peptide maintains an exchange-amenable pool of HLA class II within endo-lysosomal compartments (Cresswell, 1994), in the HLA-I pathway, natural variations in assembly and resulting stability create a pool of exchange-amenable HLA-I molecules, but in an allotype selective manner. Under steady state conditions, endo-lysosomal HLA-I assembly is likely to be important for maintaining peripheral tolerance against proteins predominantly localized to endo-lysosomes and secreted factors internalized via bulk endocytosis. Allotypes such as B*08:01 with reduced capability for endo-lysosomal assembly may be more likely to break peripheral tolerance, which could explain some known associations with autoimmune diseases (Price et al., 1999, Candore et al., 2002, Miller et al., 2015, Rothwell et al., 2016). Under inflammatory conditions, increased cross-presentation efficiency could lead to increased priming and better activation of antigen-specific CTL responses mediated by allotypes such as B*35:01. Indeed, these findings could explain why, in HIV infections, tapasin-independent allotypes such as B*35:01 have increased breadth of peptide presentation to HIV-specific T cells (Bashirova et al., 2020). Additionally, allotypes such as B*35:01 may mediate better protection against pathogens that persist within a sub-cellular endo-lysosomal niche, including *Mycobacterium tuberculosis, Toxoplasma gondii* and *Legionella pneumophila*. More studies are needed to understand the prevalence, extent, and consequences of endo-lysosomal assembly variations among HLA-I allotypes.

## MATERIALS AND METHODS

### Human sample processing and PBMC isolation

Whole blood was collected in anticoagulant citrate dextrose (ACD) tubes from healthy human donors. These donors were consented as part of an IRB-approved study (HUM00071750) and were previously genotyped at the HLA locus (Yarzabek et al., 2018). Alternatively, non-genotyped donor blood was obtained from the University of Michigan Platelet Core. About 27 mL of blood was collected in ACD tubes from each donor. For peripheral blood mononuclear cell (PBMC) isolation, blood was diluted to 50 mL with PBS + 2% FBS (PBS/FBS) and 25 mL of diluted blood was overlaid on top of 15 mL Ficoll-paque in two tubes. The tubes were spun at 400 x g for 30 minutes at room temperature in a swinging bucket rotor, with the acceleration and deceleration settings set to 4 and 0, respectively. After centrifugation, the top layer of plasma was discarded, and the center layer of cells was collected. PBMCs were washed twice with PBS/FBS for 10 min at 2500 rpm, and PBMCs were resuspended in R10 medium (RPMI + 10% FBS + 1% Antibiotic/Antimycotic + 1% L-glutamine) and counted.

### Monocyte isolation and moDC generation

For monocyte isolation, cells were purified directly from whole blood or frozen PBMCs using negative magnetic selection. Whole blood was processed using the StemCell EasySep Direct Monocyte Isolation kit (catalog # 19669) for whole blood, or the Miltenyi Classical Monocyte Isolation Kit (catalog # 130-117-337) according to the manufacturer’s instructions. After isolation, cells were washed with PBS + 1 mM EDTA (PBS/EDTA), counted, and resuspended in R10 medium at a concentration of 1 million/mL. For moDC differentiation, 6 mL cells (6 million) were added to a well of a 6-well plate, and GM-CSF and IL-4 were added to concentrations of 10 ng/mL and 50 ng/mL respectively. The top 3 mL of medium was replaced with fresh medium + IL-4 and GM-CSF on day 3 and day 5. moDCs were collected for use on day 7.

### HLA-B expression measurements

PBMCs were isolated as described above. The following antibody cocktail was diluted in PBS/FBS and used to stain PBMCs for 30 min on ice to identify various cell populations: anti-CD3-Pacific Blue, anti-CD33-APC/Cy7, anti-CD14-AF700, and anti-HLA-DR-BV650 (all used at 1:200 and from Biolegend). Monocytes were identified as FSC^int^, SSC^low^, CD3^-^, CD14^+^, CD33^+^, HLA-DR^+^. For live cell staining and surface HLA-B measurements of PBMCs, cells were aliquoted into a 96-well plate and washed with PBS, followed by staining with 100 µL of antibody cocktail + 1:40 anti-Bw6-FITC (OneLambda). For moDC identification after monocyte isolation and differentiation, cells were stained with an antibody cocktail of anti-CD11c-PE/Cy7, anti-HLA-DR-BV650, and anti-CD209-APC (all used at 1:200 and from Biolegend). After staining for 30 min on ice, cells were washed twice with PBS, then stained for 15 min at room temp with 7-AAD (1:200, BD), followed by analysis on a flow cytometer. For inhibitor treatments, PBMCs or moDCs were treated in a 96-well plate with either bafilomycin A1 (200 nM, Cayman Chemical catalog #11038) or MG132 (10 µg/mL, Sigma catalog # 474787) for various time points, and the above staining protocol was followed.

For surface and total HLA-B or HLA-C measurements of moDCs, cells were first stained with Red Fixable Live/Dead dye diluted in PBS for 15 min at room temp (1:1000, ThermoFisher). Cells were washed with PBS, followed by fixation with 4% PFA diluted in PBS for 10 min at room temp. Next, cells were washed with PBS, and half of the samples stained with the antibody cocktail + anti-Bw6 (surface HLA-Bw6), and half stained with the antibody cocktail alone. Cells were stained for 30 min on ice, then washed twice. The cells stained for surface HLA-Bw6 were set aside, and the remaining cells were stained with anti-Bw6 diluted in 0.2% saponin (total HLA-Bw6) for 30 min on ice. After washing twice, both sets of cells were analyzed on a BD LSR Fortessa flow cytometer. Data was analyzed using FlowJo software.

### moDC HLA-B half-life measurements

Surface stability and half-life assessment was performed as described previously(Zarling et al., 2003, Yarzabek et al., 2018). Briefly, monocytes were isolated and differentiated to moDCs for 7 days as described above. B*08:01^+^ or B*35:01^+^ donor moDCs were plated into a 96-well plate in duplicate for each condition. Brefeldin A (BFA) treatment was added at negative time-points: for a four-hour time course, BFA was first added to cells for the hour four treatment time point, then hour three, etc. BFA was added at a concentration of 0.5 µg/mL to each well in media, and cells were incubated before centrifugation, washing with PBS, and staining with a monoclonal antibody cocktail of anti-CD11c-PE/Cy7, anti-HLA-DR-BV650, and anti-CD209-APC (all used at 1:200 and from Biolegend), as well as anti-Bw6-FITC (Biolegend, 1:40), for 30 min on ice. After staining, cells were washed twice with PBS, followed by staining with 7-AAD and analysis on a BD LSR Fortessa flow cytometer. Half-life values were extracted using a one phase decay curve with a constrained plateau of zero.

### moDC peptide receptivity

Peptides were synthesized by A&A Labs LLC. Monocytes were isolated and differentiated to moDCs for 7 days as described above. B*08:01^+^ or B*35:01^+^ donor moDCs were plated into a 96-well plate in duplicate for each condition, followed by the addition of either DMSO, canonical peptide (100 µM), or control peptide (100 µM). Control peptides were truncated and altered at anchor residues relative to the canonical peptide sequence. For B*08:01, the canonical peptides used were HSKKKCDEL or GPKVKRPPI, and the control peptides used were HSDYECDE or GPDVERPP. For B*35:01, the canonical peptides used were HPVGEADYFEY or EPLPQGQLTAY, and the control peptides used were HGVGEADYFE or EGLPQGQLTA. Peptides were incubated with moDCs for 4 hrs at 37°C, followed by washing and staining with antibodies for moDC surface markers, as well as the monoclonal antibody HC10-FITC, which recognizes peptide-deficient conformers of HLA-I. After 30 min of staining on ice, cells were washed twice with PBS, stained with 7-AAD (1:200), and analyzed on a BD LSR Fortessa flow cytometer. Additional experiments were performed where peptide incubation was performed at 4°C, or at 37°C in the presence of 200 nM bafilomycin. Data was analyzed using FlowJo software.

### Confocal microscopy

Monocytes were isolated from blood as described above. Glass coverslips were coated with poly-L-lysine for 2 hrs at 37°C in 12-well plates, then washed 3X with water and allowed to dry completely. For each coverslip, about 250,000 monocytes were added in 100 µL medium, and allowed to adhere to the coverslips for 2 hrs at 37°C. Coverslips were washed with PBS gently, then fixed with 4% PFA for 10 min at room temp. Coverslips were washed with PBS, then permeabilized with 0.1% Triton X-100 in PBS for 10 min at room temp. Coverslips were washed twice with PBS, then blocked with 5% goat serum diluted in PBS + 0.05% Tween 20 (PBST). Blocking was performed for 1 hr at room temp with gentle rocking. After blocking, coverslips were inverted onto a 100 µL bubble of primary antibody staining solution (diluted antibody + 1% BSA in PBST) placed on a piece of parafilm and incubated in a cold room overnight. The next day, the coverslips were returned to a 12-well plate and washed 3X with PBST for 5 min each with rocking. Coverslips were stained with 500 µL secondary antibody solution (antibody diluted in PBST + 1% BSA) for 1 hr at room temp while rocking. Coverslips were washed 3X with PBST for 5 min each with rocking and inverted onto a 15 µL drop of ProLong Diamond + DAPI placed on a glass slide. Slides were cured at room temp overnight, then sealed with nail polish around the edges. Images were acquired with a Nikon A1 confocal microscope using a pinhole size of 1, a z-step of 0.3 um, a pixel dwell time of 12.1, and a line average of 2. Primary antibodies used were: mouse anti-Bw6-biotin (1:20, OneLambda), rabbit anti-EEA1 (1:1000, Invitrogen), mouse IgG1 anti-Arf6 (1:20, Invitrogen), rabbit anti-Rab11a (1:12.5, ThermoFisher), rabbit anti-LAMP1 (1:100, CellSignaling Technologies). Secondary antibodies/probes used were: streptavidin-AF488 (1:2000, Invitrogen), streptavidin-AF647 (1:2000, Invitrogen), goat anti-rabbit-AF555 (1:500, Abcam), goat anti-mouse IgG1-AF488 (1:500, Invitrogen).

For confocal experiments using bafilomycin treatment, moDCs were cultured and allowed to adhere to coverslips as described above. After adherence for 1-2 hrs, culture media in each well was replaced with either 1 mL R10 for the untreated controls, or 1 mL R10 + 200 nM bafilomycin for treatment conditions. Cells were cultured with inhibitor for 4 hrs, followed by washing with PBS, fixation for 10 min with 4% PFA, and staining for Bw6/Rab11a or Bw6/LAMP1 as described above.

Co-localization was assessed using one of two methods. For object-based co-localization, a FIJI macro was written based on a previously described method (Moser et al., 2017). Briefly, image files were analyzed using an identical macro script involving background masking and subtraction, signal thresholding, and quantification of the fraction of signal A that overlaps spatially with signal B. Pearson’s correlation was performed uniformly to each image file using the JACOP plugin for FIJI (Bolte and Cordelières, 2006), which quantifies the correlation of co-occurrence of bright pixels of signal A and bright pixels of signal B.

### Antigen uptake time course

Monocytes or moDCs were plated into a 96-well plate at 100,000 cells/well. Cells were pulsed with 10 µg/mL BSA labeled in-house with Alexa Fluor 594 in duplicate for either 15 min, 30 min, 45 min, or 60 min. Duplicate wells were left untreated, and duplicate wells were pulsed with BSA-AF594 for 60 min on ice to inhibit endocytosis and measure background fluorescence. After each time point, cells were collected and washed twice with PBS, followed by fixation with 4% PFA for 5 min at room temp. Cells were washed and analyzed by flow cytometry using a BD LSR Fortessa. Data was analyzed using FlowJo software.

### DQ-Ova antigen processing assays

Monocytes or moDCs were plated into a 96-well plate at 100,000 cells/well. Cells were pulsed with 50 µg/mL DQ-Ova antigen in media for 30 min at either 37°C or 4°C. Following 30 min pulse, cells were washed with PBS and chased in either media alone, media + 200 nM bafilomycin, media + 10 µg/mL MG132, or media + bafilomycin and MG132 for 2 hrs (monocytes) or 4 hrs (moDCs). Control cells which were pulsed with DQ-Ova at 4°C were also chased in media at 4°C for 2 or 4 hrs. Following chase, cells were washed with PBS and stained with Aqua fixable live/dead (1:500 in PBS, ThermoFisher catalog #L34957) for 15 min at room temp. Cells were washed again with PBS and fixed with 4% PFA for 10 min at room temp. Samples were measured by flow cytometry using a BD LSR Fortessa, and analyzed using FlowJo software.

### Polyclonal antigen-specific CTL expansion

CD8^+^ T cells were isolated by negative magnetic selection from either B*08:01^+^ or B*35:01^+^ donor blood using kits from StemCell Technologies (catalog #17953). To screen for antigen-specific CTLs, common EBV epitopes for a particular HLA-B allele were identified from the Immune Epitope Database (IEDB) and used to produce peptides. These peptides were loaded onto B*08:01 or B*35:01 monomers as described above, and peptide-loaded monomers were bound to streptavidin-APC molecules for flow cytometric staining. Isolated CD8^+^ T cells were stained with tetramer (1:20) for 1 hr on ice. Cells were washed with PBS and stained with anti-CD3-Pacific Blue (1:200, Biolegend catalog #300417) and anti-CD8-AF700 (1:200, Biolegend catalog #344724) antibodies diluted in PBS/FBS for 30 min on ice. Cells were washed twice and stained with 7-AAD (1:200) and analyzed by flow cytometry. Tetramer positive cells were identified and sorted via fluorescence-activated cell sorting (FACS).

Sorted tetramer-specific CTLs were expanded as previously described (Dong et al., 2010). Briefly, HLA-I allo PBMCs were isolated and resuspended to a concentration of 2 million cells/mL. Cells were irradiated at 3300 rad (performed at the UMich Experimental Irradiation Core) and plated at 200,000 cells/well in a 96-well plate with 3.2 µg/mL phytohemagglutinin (PHA, Remel catalog # R30852801). About 1000 sorted cells were added to each well, to a final volume of 200 µL/well. Twice per week, the top 100 µL of media was removed from each well and fresh media + 10 µL natural human IL-2 (Hemagen Diagnostics, catalog # 906011) was added. CTLs were maintained in this feeder cell expansion culture for 2-3 weeks, checking the expansion of tetramer^+^ cells by the end with of the expansion. Following expansion in 96-well plate format, CTLs were transferred to a T-25 flask with 25-50 million irradiated allo-PBMCs, 1 µg/mL PHA, and 50 U/mL recombinant human IL-2 (Peprotech, catalog # 200-02) in 25 mL media. Expansion in flasks was continued for another 2-3 weeks, checking with tetramer staining periodically. CTL density was kept under 2 million cells/mL, splitting into a new flask as needed during the expansion. After expansion, CTLs were frozen down into aliquots of about 5 million cells per cryovial. For use in activation assays, a CTL vial was thawed and added to a T-25 flask with 25-50 million irradiated allo-mismatched PBMCs, PHA, and IL-2. CTLs typically re-expanded to a usable density in about 1-1.5 weeks, and were ready to use for activation assays for the next 1-2 months.

### CTL peptide titration

PBMCs from B*08:01^+^ or B*35:01^+^ were pulsed overnight with media alone, or media + serially-diluted peptides. Peptide concentrations ranged from 1 pM to 100 µM, with 10-fold dilutions. PBMCs pulsed in duplicate for each condition, at 100,000 cells/well in a 96-well plate. After overnight incubation with peptide, PBMCs were washed and co-cultured with CTLs specific for each peptide at a 2:1 CTL:PBMC ratio (200,000 CTLs/well). GolgiStop and GolgiPlug inhibitors (1:800) were included in the co-culture to block cytokine export, and an anti-CD107a-PE antibody (1:200) was added to stain for CTL degranulation. After a 5-hour co-culture, cells were washed and stained with Red Fixable Live/Dead dye (1:1000) for 15 min at room temp. Cells were washed, then stained with anti-CD3-Pacific Blue (1:200) and anti-CD8-AF700 (1:200) antibodies diluted in PBS/FBS for 30 min on ice. Cells were washed twice and fixed for 10 min at room temp with 4% PFA diluted in PBS. After fixation, cells were stained intracellularly with anti-IFNγ-FITC (1:100, Biolegend catalog #506504) diluted in 0.2% saponin/PBS for 30 min on ice. Cells were washed twice and analyzed by flow cytometry using a BD LSR Fortessa. Data was analyzed by FlowJo.

### Plasmid cloning

A BZLF1 encoding sequence was synthesized by Integrated DNA Technologies (IDT) based on the European Nucleotide Archive coding sequence (accession # AAA66529). Primers were used to perform PCR to amplify the sequence:

5’-TAAGCAGGATCCATGATGGACCCAAACTCGAC-3’ and

5’-TGCTTAGCGGCCGCTTAGAAATTTAAGAGATCCT-3’.

The primers inserted a BamHI site upstream of the BZLF1 gene, and a NotI site downstream of the gene, with about 6 nucleotides of overhang on either side of the PCR product. PCR product was digested with BamHI and NotI, after which the enzymes were removed using a PCR Cleanup Kit (Qiagen). The vector pGEX-4T-LP was digested using the same enzymes, ran on a 1% agarose gel, excised, and cleaned up using a Gel Extraction Kit (Qiagen). The BZLF1 PCR product was ligated into the pGEX vector using T4 DNA ligase at 16°C overnight. The ligation product was transformed into Rosetta cells, which were selected on LB-Ampicillin plates. Colonies were picked, plasmid DNA isolated, and sequenced to confirm gene insertion. The BamHI site places the BZLF1 gene in-frame downstream of the GST protein, creating a GST-BZLF1 fusion protein containing a thrombin cleavage site between the proteins.

The EBNA3A:133-491 (simplified to EBNA3A throughout) truncation protein was cloned by PCR amplification of a segment of the gene encoding residues 133-491. This segment contained the coding sequence for both the FLRGRAYGL and YPLHEQHGM epitopes. The MSCV-N EBNA3A plasmid used as a backbone for PCR was a gift from Karl Munger (Addgene plasmid # 37956; http://n2t.net/addgene:37956.; RRID:Addgene_37956) (Rozenblatt-Rosen et al., 2012). Primers used to amplify the EBNA3A:133-491 gene segment: 5’-CGCGTGGATCCATGTACATAATGTATGCCATGGC -3’ and

5’-ACGATGCGGCCGCTTAAACACCTGGGAGTTG -3’.

The primers inserted a BamHI site upstream of the EBNA3A:133-491 gene, and a NotI site downstream of the gene, with about 6 nucleotides of overhang on either side of the PCR product. Cloning proceeded as described for BZLF1 above.

### Protein expression and purification

GST-BZLF1 protein or GST-EBNA3A:133-491 protein was purified based on the protocol by Harper and Speicher (Harper and Speicher, 2011). Briefly, Rosetta cells were transformed with either BZLF1-pGEX-4T-LP or EBNA3A-pGEX-4T-LP plasmids and plated on LB-Amp plates, and single colonies were picked and used to inoculate a 30 mL LB-Amp overnight starter culture. The next day, 10 mL of culture was used to inoculate a 1 L flask of LB-Amp. The flask was grown shaking at 37°C until the culture reached an OD600 of between 0.5 and 0.7. The flask was then cooled to 16°C and IPTG was added to a final concentration of 1 mM. Protein expression was induced overnight at 16°C with shaking. The next day, the culture was split in half and spun down for 30 min at 4000 g at 4°C. One pellet was stored at -80°C, and the other was resuspended in 15 mL lysis buffer (50 mM Tris + 1% Triton X-100 + 5 mM EDTA + 1 mM 2-ME + 0.15 mM PMSF + 1 cOmplete protease inhibitor tablet/50 mL, pH = 8.0) and lysed by sonication. The lysate was spun at 13000 rpm at 4°C for 30 min, and the supernatant was applied to glutathione resin. The supernatant and column were incubated gently rocking at 4°C for 2 hrs. The column was thoroughly washed with 100 mL PBS + 5 mM EDTA + 0.15 mM PMSF (PBS/EDTA/PMSF), followed by 100 mL PBS/EDTA. 20 mL of 20 mM reduced L-glutathione was applied to the column and allowed to sit at room temp for 30 min. The glutathione solution was allowed to slowly move through the column and eluted protein was collected in 1 mL fractions. Eluted fractions were quantified for protein concentration using a nanodrop, and selected fractions were analyzed using 12% SDS PAGE gel alongside lysate fractions to check protein purity and size. Fractions containing the GST-BZLF1 or GST-EBNA3A fusion were concentrated using a 10 k MWCO centricon and buffer exchanged into PBS buffer. Protein was concentrated typically to a concentration of about 2 mg/mL, and stored at -20°C.

### Cross-presentation assays

Monocytes or moDCs were plated into sterile 96-well plates at 50,000-100,000 cells/well. In quadruplicate, APCs were pulsed with either no antigen, peptide (50 µM), 100 µg GST-BZLF1, or 100 µg GST-EBNA3A. For the inhibitor experiments, cells were pulsed with 100 µg protein (GST-BZLF1 or GST-EBNA3A), 100 µg protein + 50 µM CTI1 (Selleck Chem catalog # S2847), or 100 µg protein + 10 µg/mL MG132 (Sigma catalog # 474787). Antigen pulses with and without inhibitors were performed for 6 hr at 37°C, followed by washing with PBS. The two antigen-specific CTLs used for comparison were added at a 1:1 CTL:APC ratio so that each CTL was added to each condition in duplicate. Also added to the culture with the CTLs was anti-CD107a-PE (1:20, BD, or 1:200, Biolegend), GolgiStop (1:800, BD), and GolgiPlug (1:800, BD). Cells were spun for 3 min at 1200 rpm, and co-incubated for 5 hr. After co-culture, cells were washed and stained with Red Fixable Live/Dead dye (1:1000, ThermoFisher catalog #L34971) for 15 min at room temp. Cells were washed, then stained with anti-CD3-Pacific Blue (1:200, Biolegend catalog #300417) and anti-CD8-AF700 (1:200, Biolegend catalog #344724) antibodies diluted in PBS/FBS for 30 min on ice. Cells were washed twice and fixed for 10 min at room temp with 4% PFA diluted in PBS. After fixation, cells were stained intracellularly with anti-IFNγ-FITC (1:100, Biolegend catalog #506504) diluted in 0.2% saponin/PBS for 30 min on ice. Cells were washed twice and analyzed by flow cytometry using a BD LSR Fortessa. Data was analyzed by FlowJo.

## List of Supplementary Materials

**Table S1. Healthy human donors and HLA genotypes used in study**.

**Figure S1: Human moDCs express HLA-B at least 4 times higher than HLA-C on the cell surface Figure S2. Human moDCs have lower HLA-Bw6 cell surface stability compared to monocytes and lymphocytes**.

**Figure S3. Pearson’s correlation analysis of monocyte and moDC endo-lysosomal co-localization**.

**Figure S4. Reduced B*35:01 dependence on proteasomal processing in monocytes and moDCs, and additional HLA-Bw6 co-localization studies in bafilomycin-treated cells**.

**Figure S5. Peptide and soluble antigen presentation by monocytes and moDCs**.

## Acknowledgements

We thank all of our study participants for their time and participation in the study, as well as the Michigan Clinical Research Unit (MCRU) for performing blood draws. We thank Kole Tison for the assistance with the study. We thank the NIH Tetramer Core for the B*08:01 and B*35:01 tetramers used for the isolation of antigen-specific CD8^+^ T cells. We thank Aaron Taylor and Eric Rentchler at the Microscopy Core of the University of Michigan Biomedical Research Core Facility and Miles Mckenna (Nikon Instruments Inc.) for assistance with user training or microscopy image analysis. We thank Amanda Prieur and the UM Platelet Core for providing non-genotyped blood for experiments. We also thank staff at the Flow Cytometry Core the University of Michigan Biomedical Research Core Facility for assistance with cell sorting and Dr. David Karnak and the Experimental Irradiation Core at Michigan for assistance producing irradiated feeder cells for CTL expansion.

