## Supplemental Table and Figures for "Endo-lysosomal assembly variations among Human Leukocyte Antigen class I (HLA-I) allotypes"

**Supplementary Materials:**

| GROUP 1 | B*08:01$^{+}$ donors: either B*08:01 double positive, or B*08:01/Bw4 heterozygous | | | | | | |
|---|---|---|---|---|---|---|---|
| | **Donor ID** | **HLA-A** | **HLA-A** | **HLA-B (Bw6)** | **HLA-B (Bw4 or B*08:01)** | **HLA-C** | **HLA-C** |
| | 9 | A*01:01:01:01 | A*01:01:01:01 | B*08:01:01 | B*08:01:01 | C*07:01:01:01 | C*07:01:01:01 |
| | 28 | A*01:01:01:01 | A*02:01:01:01 | B*08:01:01 | B*51:01:01:01 | C*07:01:01:01 | C*15:13 |
| | 55 | A*01:01:01:01 | A*23:01:01 | B*08:01:01 | B*44:03:01 | C*07:01:01:01 | C*04:09 |
| | 94 | A*01:01:01:01 | A*68:01:02:01 | B*08:01:01 | B*44:02:01:01 | C*07:01:01:01 | C*05:01:01:02 |
| | 121 | A*01:01:01:01 | A*01:01:01:01 | B*08:01:01 | B*27:05:02 | C*07:01:01:01 | C*02:07 |
| | 130 | A*01:01:01:01 | A*30:01:01 | B*08:01:01 | B*13:02:01 | C*07:01:01:01 | C*06:02:01:01 |
| | 137 | A*01:01:01:01 | A*01:01:01:01 | B*08:01:01 | B*37:01:01 | C*07:01:01:01 | C*06:02:01:01 |
| | 166 | A*01:01:01:01 | A*01:01:01:01 | B*08:01:01 | B*08:01:01 | C*07:01:01:01 | C*07:01:01:01 |
| | 178 | A*01:01:01:01 | A*02:01:01:01 | B*08:01:01 | B*57:01:01 | C*07:01:01:01 | C*06:02:01:01 |
| | 237 | A*68:01:01:02 | A*02:01:01:01 | B*08:01:01 | B*44:02:01:01 | C*07:01:01:01 | C*05:01:01:02 |
| **GROUP 2** | **B*35:01$^{+}$ donors: either B*35:01 double positive, or B*35:01/Bw4 heterozygous** | | | | | | |
| | **Donor ID** | **HLA-A** | **HLA-A** | **HLA-B (Bw6)** | **HLA-B (Bw4)** | **HLA-C** | **HLA-C** |
| | 24 | A*02:01:01:01 | A*24:02:01:01 | B*35:01:01:02 | B*51:01:01:01 | C*15:02:01:01 | C*04:04:01 |
| | 136 | A*11:01:01:01 | A*30:01:01 | B*35:01:01:02 | B*13:02:01 | C*04:01:01:01 | C*06:02:01:01 |
| | 141 | A*02:01:01:01 | A*03:01:01:01 | B*35:01:01:02 | B*44:02:01:01 | C*05:01:01:02 | C*04:01:01:01 |
| | 168 | A*02:01:01:01 | A*11:01:01:01 | B*35:01:01:02 | B*51:01:01:01 | C*15:02:01:01 | C*04:01:01:01 |
| | 187 | A*01:01:01:01 | A*02:01:01:01 | B*35:01:01:02 | B*44:02:01:01 | C*05:01:01:02 | C*04:01:01:05 |
| | 210 | A*01:01:01:01 | A*03:01:01:01 | B*35:01:01:02 | B*57:01:01 | C*06:02:01:01 | C*04:01:01:01 |
| **GROUP 3** | **B*08:01$^{+}$ or B*35:01$^{+}$ donors for antigen-specific CTL expansion** | | | | | | |
| | **Donor ID** | **HLA-A** | **HLA-A** | **HLA-B** | **HLA-B** | **HLA-C** | **HLA-C** |
| | 9 | A*01:01:01:01 | A*01:01:01:01 | B*08:01:01 | B*08:01:01 | C*07:01:01:01 | C*07:01:01:01 |
| | 16 | A*01:01:01:01 | A*02:01:01:01 | B*08:01:01 | B*35:01:01:02 | C*07:01:01:01 | C*04:01:01:06 |
| | 40 | A*03:01:01:01 | A*11:01:01:01 | B*35:01:01:02 | B*35:03:01 | C*04:01:01:01 | C*04:01:01:01 |
| **GROUP 4** | **B*08:01/B*35:01 double positive donors for cross-presentation APCs** | | | | | | |
| | **Donor ID** | **HLA-A** | **HLA-A** | **HLA-B** | **HLA-B** | **HLA-C** | **HLA-C** |
| | 16 | A*01:01:01:01 | A*02:01:01:01 | B*08:01:01 | B*35:01:01:02 | C*07:01:01:01 | C*04:01:01:06 |
| | 25 | A*01:01:01:01 | A*02:01:01:01 | B*08:01:01 | B*35:01:01:02 | C*07:01:01:01 | C*04:01:01:06 |
| | 132 | A*01:01:01:01 | A*24:02:01:01 | B*08:01:01 | B*35:01:01:02 | C*07:01:01:01 | C*07:01:01:01 |

**Table S1. Healthy human donors and HLA genotypes used in study.**

Donors were selected from our previously described cohort of HLA genotyped healthy participants (Yarzabek et al. 2018). Four primary groups were recruited for this study. Group 1 donors were either homozygous for B*08:01 at the HLA-B locus or had Bw6/Bw4 heterozygosity at the HLA-B locus with B*08:01 as the Bw6 allele. Group 2 donors had Bw6/Bw4 heterozygosity at the HLA-B locus with B*35:01 as the Bw6 allele. Group 3 donors expressed B*08:01, B*35:01, or both. Heterozygosity for Bw6/Bw4 was not required for studies with this group. Group 4 contained donors heterozygous for

B*08:01 and B*35:01. Groups 1 and 2 were used to measure the expression and localization of B*08:01 and B*35:01 respectively with an anti-Bw6 monoclonal antibody. HLA-C's cross-reactive with anti-Bw6 are indicated in blue or red. No HLA-A's are cross reactive with anti-Bw6 (Yarzabek et al. 2018). Note that the HLA-B*08:01 homozygous cells are the only ones with two cross-reactive HLA-C*07:01. Group 3 was used as effector antigen-specific CTLs for all T cell activation assays. Group 4 was used as antigen-presenting cells for cross-presentation assays. For assays where specific HLA genotype was not required, such as DQ-Ova experiments, non-genotyped donors 248-275 were used, or blood was obtained from the University of Michigan Platelet Core. Donors from the Platelet Core are labeled with the prefix "PCD".

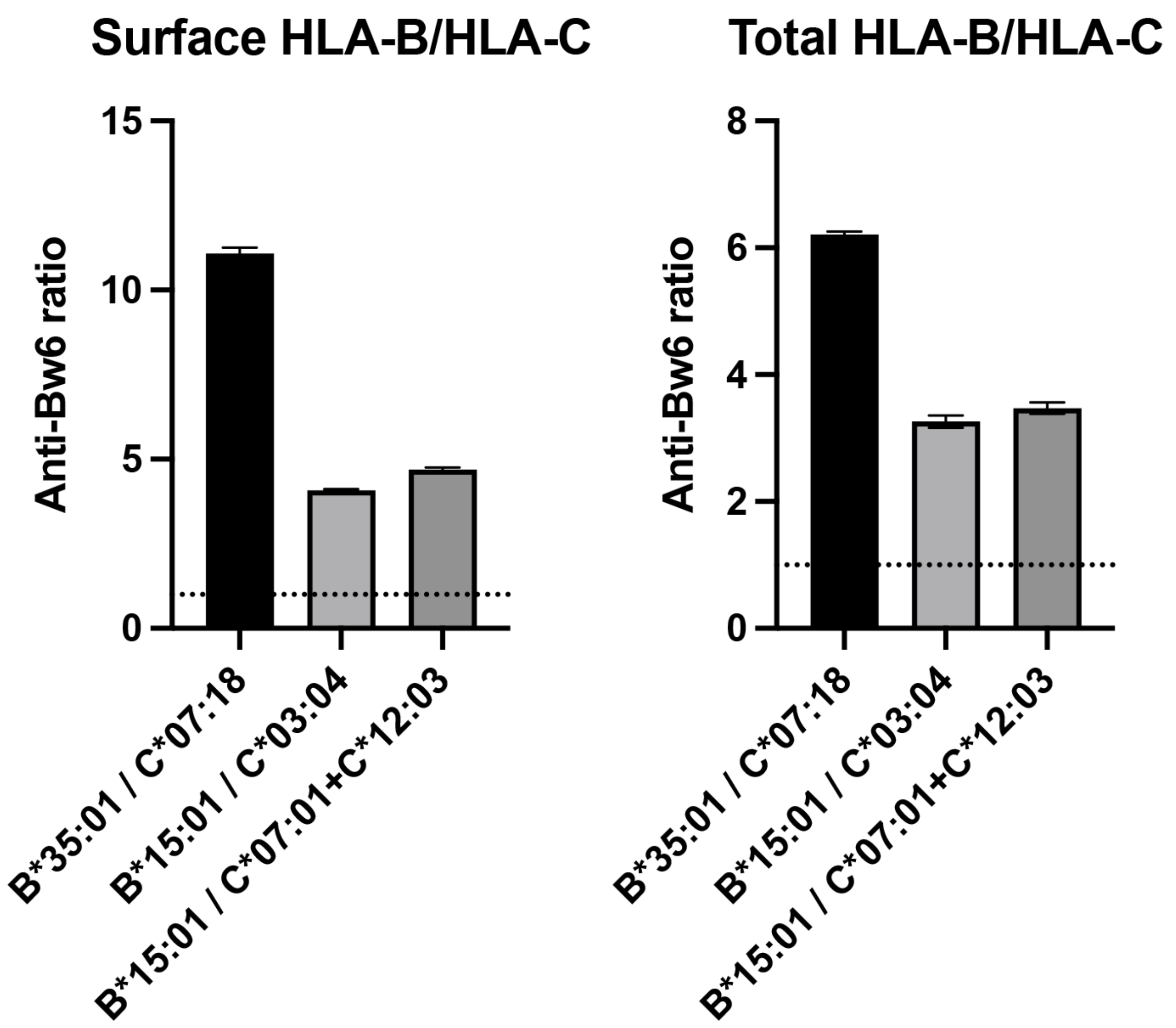

**Figure S1. Human moDCs express HLA-B at least 4 times higher than HLA-C on the cell surface.**

Donors were selected expressing either a single HLA-B allotype with a Bw6 epitope and no cross-reactive HLA-C (called HLA-B donor), or those with no anti-Bw6-reactive HLA-B, but with one to two HLA-C's containing a Bw6 epitope (called HLA-C donor). For each experiment, an HLA-B donor was recruited with an HLA-C donor for parallel measurements of cell-derived anti-Bw6 MFI values. Monocytes were isolated and differentiated to moDCs, followed by fixation and staining for surface Bw6, or fixation, permeabilization, and staining for total Bw6. The anti Bw6 MFI values from the HLA-B donors were divided by the anti-Bw6 signals from the HLA-C donors to find the fold difference in HLA-B expression relative to HLA-C expression, which ranged from 3- to 11-fold. HLA-B donors were: 24 (A*02:01, A*24:02, **B*35:01**, B*51:01, C*15:02, C*04:04) and 124 n=2 (A*32:01, A*11:01, **B*15:01**, B*53:01, C*06:02, C*04:01). HLA-C donors were: 205 (A*01:01, A*68:01, B*13:02, B*58:01, **C*07:18**, C*06:02), 38 (A*02:01, A*02:01, B*13:02, B*13:01, **C*03:04**, C*06:02), and 76 (A*24:02, A*01:01, B*15:17, B*38:01, **C*07:01**, **C*12:03**).

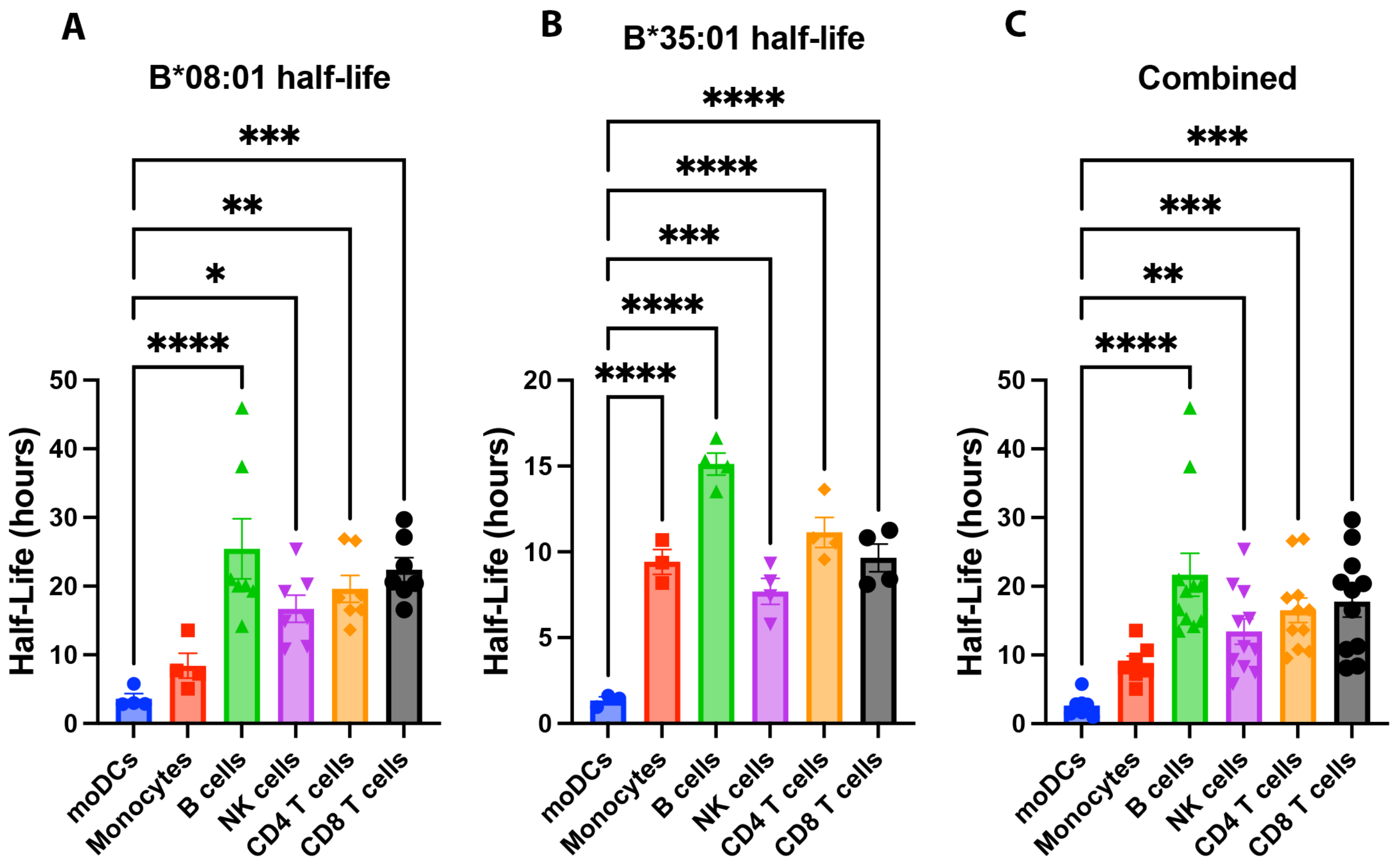

**Figure S2. Human moDCs have lower HLA-Bw6 cell surface stability compared to monocytes and lymphocytes.** moDC Bw6 half-life data from B*08:01⁺ **(A)** B*35:01⁺ **(B)** or the combined (B*08:01⁺ and B*35:01⁺) **(C)** donors were compared to Bw6 half-life data in monocytes or various lymphocyte populations via a One-way ANOVA analysis. Monocyte and lymphocyte data are from our previously published study (Yarzabek et al. 2018). moDC data are from Figure 1. Data repeated for donors across different experiments was averaged, so each data point represents an averaged donor half-life.

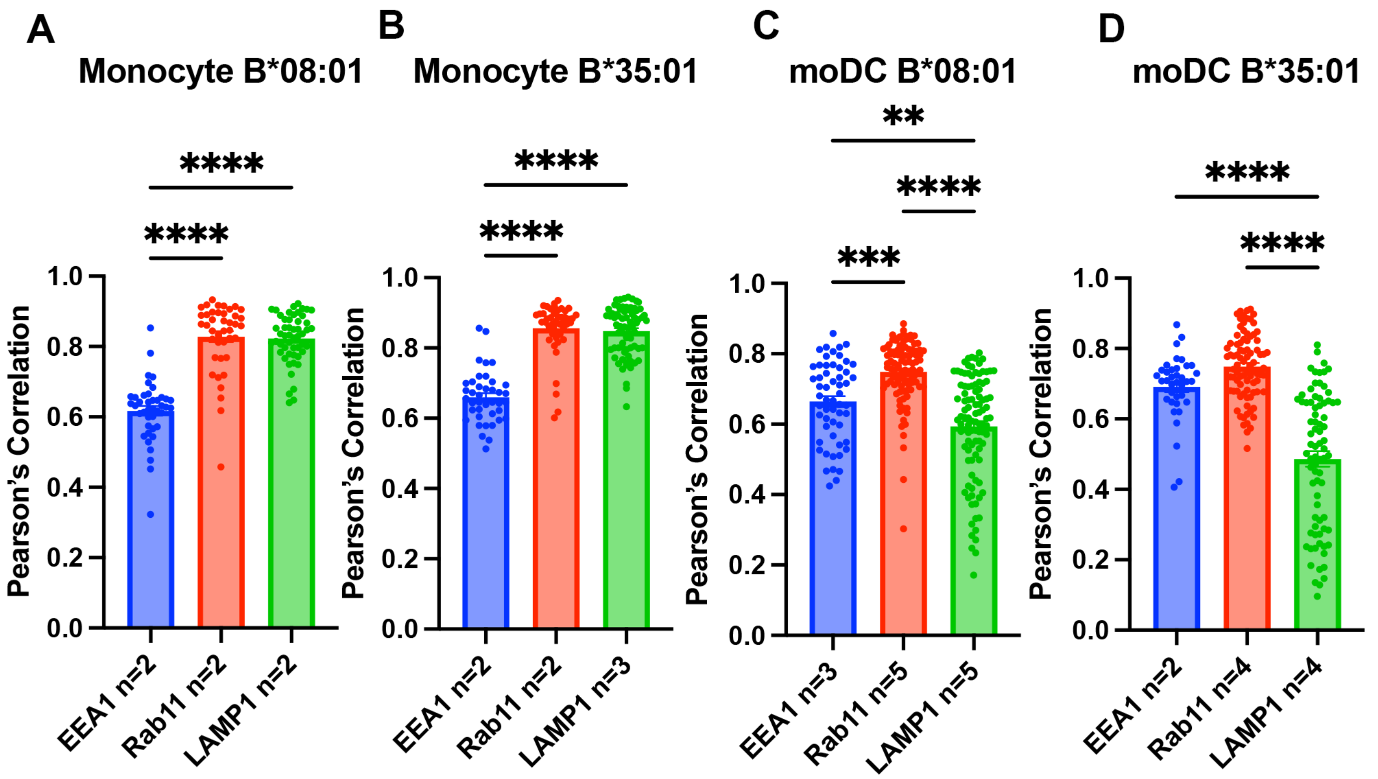

**Figure S3. Pearson's correlation analysis of monocyte and moDC endo-lysosomal co-localization.**

Data in Figure 2 were analyzed using the JACOP plugin on FIJI to calculate the Pearson's correlation between HLA-Bw6 and each of the three endo-lysosomal markers. Monocytes from B*08:01$^{+}$ **(A)** and B*35:01$^{+}$ **(B)** donors as indicated were used to measure Bw6 co-localization with EEA1, Rab11, and LAMP1. moDCs from B*08:01$^{+}$ **(C)** and B*35:01$^{+}$ **(D)** donors as indicated were used to measure Bw6 co-localization with EEA1, Rab11, and LAMP1. One-way ANOVA analyses were used to compare co-localization of HLA-Bw6 with each marker.

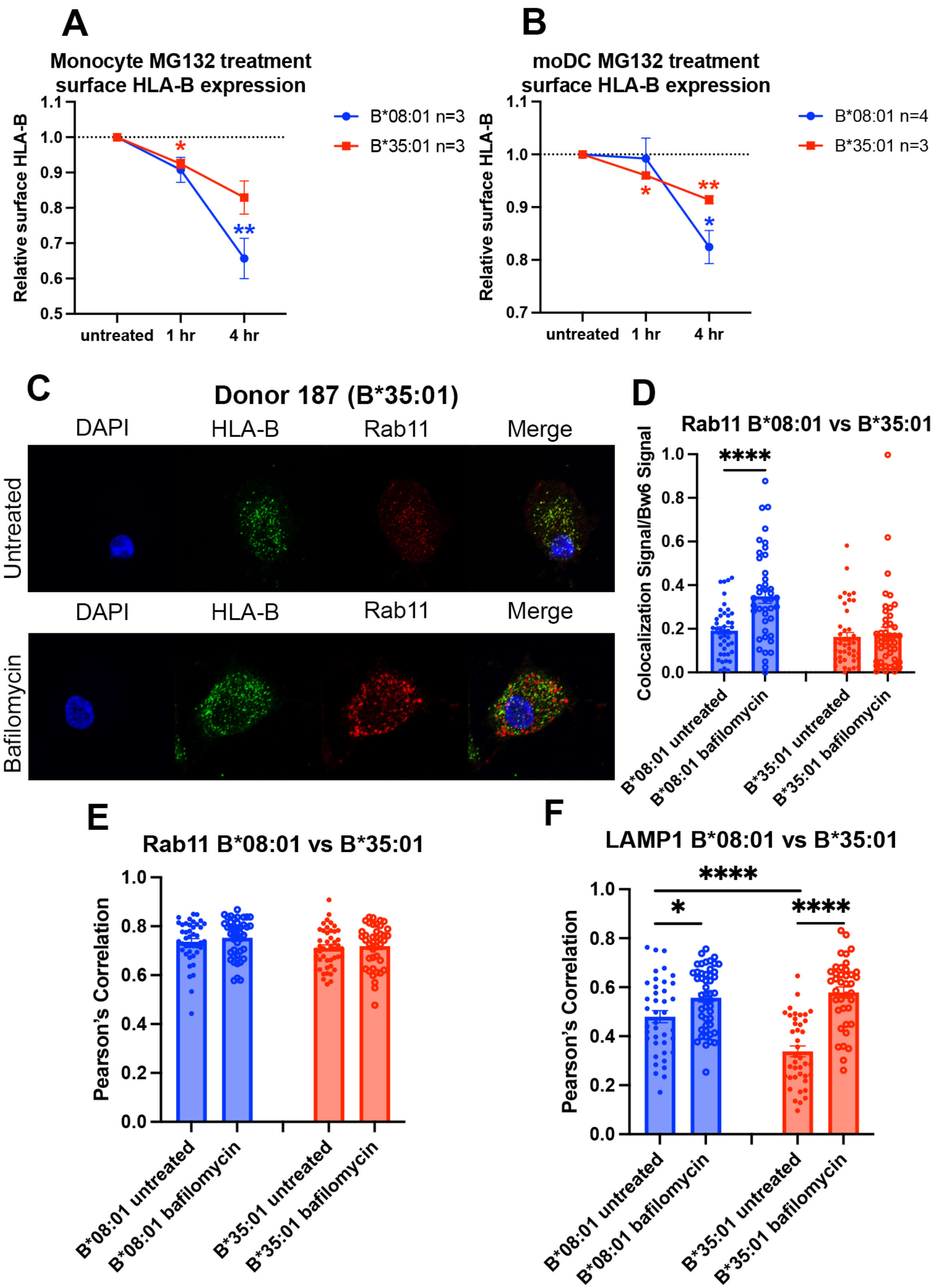

**Figure S4. Reduced B*35:01 dependence on proteasomal processing in monocytes and moDCs, and additional HLA-Bw6 co-localization studies in bafilomycin-treated cells.**

**(A)** Monocytes were treated with MG132 over a 4-hour time course, and surface expression of HLA-Bw6 measured in B*08:01⁺ and B*35:01⁺ cells. Change in expression plotted relative to untreated. B*08:01⁺ donors were: 55, 121, 137; n=3.

B*35:01$^+$ donors were: 24, 136, 210; n=3. B*08:01 and B*35:01 surface expression changes relative to untreated were assessed by one sample t tests. **(B)** moDCs were treated with MG132 over a 4-hour time course, and surface expression of B*08:01 and B*35:01 measured. Change in expression plotted relative to untreated. B*08:01 donors were: 55 (n=2), 166, 198; n=5 independent experiments. B*35:01 donors were: 24 (n=2) and 187; n=3 independent experiments. B*08:01 and B*35:01 surface expression changes relative to untreated were assessed by one sample t tests. **(C)** Representative images of moDCs with or without bafilomycin treatment stained for HLA-Bw6 and Rab11. **(D)** Object-based co-localization of HLA-Bw6 B*08:01 (n=2) or B*35:01 (n=2) with Rab11 with and without bafilomycin treatment. B*08:01$^+$ donors were 94 and 237 (n=2), and the B*35:01$^+$ donors were 168 and 187 (n=2). Unpaired t tests were used to compare co-localization with and without bafilomycin. **(E)** Pearson's correlation of B*08:01 (n=2) or B*35:01 (n=2) with Rab11 with and without bafilomycin treatment. B*08:01$^+$ donors were 94 and 237, and the B*35:01$^+$ donors were 168 and 187. Unpaired t tests were used to compare co-localization with and without bafilomycin. **(F)** Pearson's correlation of B*08:01 (n=2) or B*35:01 (n=2) with LAMP1 with and without bafilomycin treatment. B*08:01$^+$ donors were 94 and 237, and the B*35:01$^+$ donors were 168 and 187. Unpaired t tests were used to compare co-localization with and without bafilomycin.

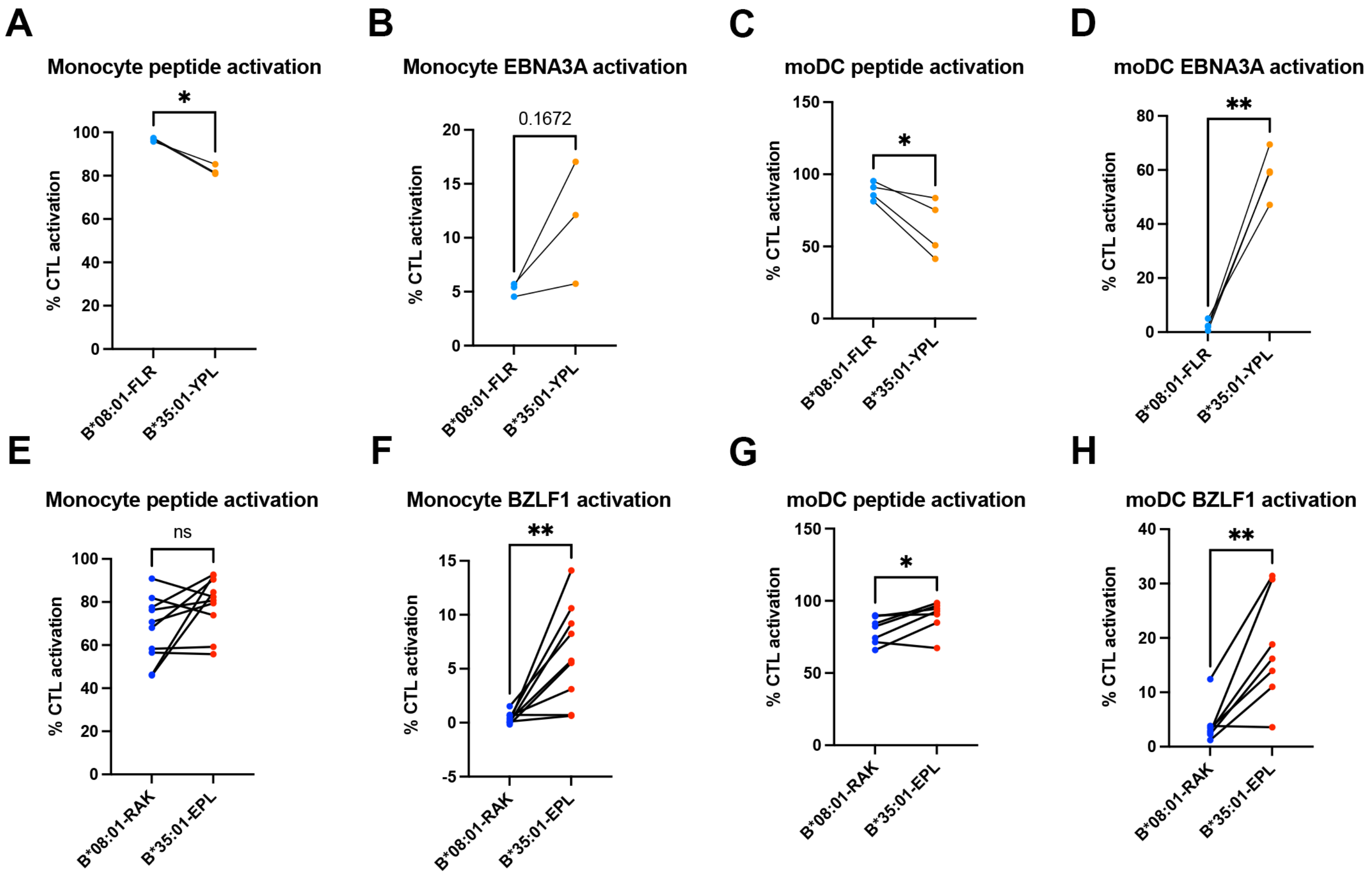

**Figure S5. Peptide and soluble antigen presentation by monocytes and moDCs.**

**(A-D)** Antigen presentation of peptide and soluble EBNA3A antigen to B*08:01-FLR and B*35:01-YPL CTLs. Monocyte peptide **(A)** and EBNA3A **(B)** presentation n=3 experiments. moDC peptide **(C)** and EBNA3A **(D)** presentation n=4 experiments. **(E-H)** Antigen presentation of peptide and soluble BZLF1 antigen to B*08:01-RAK and B*35:01-EPL CTLs. Monocyte peptide **(E)** presentation n=10 experiments, and BZLF1 **(F)** presentation n=9 experiments. moDC peptide **(G)** and BZLF1 **(H)** presentation n=7 experiments. All B*08:01 vs B*35:01 experiments shown were performed in matched experiments, and analyzed with paired t tests.
